# Maximum trophic level predicts food webs’ susceptibility to coextinctions

**DOI:** 10.64898/2026.08.14.744947

**Authors:** Henry Li, Anna Eklöf, György Barabás, Laura E. Dee

**Affiliations:** Department of Ecology and Evolutionary Biology, University of Colorado Boulder, Colorado, 80309, USA; Division of Biology, Dept. IFM, Linköping University, SE-58183 Linköping, Sweden; Institute of Evolution, Centre for Ecological Research, Budapest, Hungary

**Keywords:** coextinctions, secondary extinctions, robustness, stability, extinction risk, extinction threats, food webs, Bayesian networks

## Abstract

As ecosystems face a growing number of threats, coextinctions (resultant extinctions following a primary extinction) are expected to proliferate. However, less is known about the conditions under which coextinctions could outpace primary extinctions. Because coextinctions often occur through lost species interactions, we posit that aspects of food web structure and complexity can help predict differences in vulnerability to coextinction across ecosystems. To test this, we leverage Bayesian network models to assess the extent to which variation in ecosystem vulnerability to coextinction varies with food web structure. We find that food webs with high maximum trophic level are most vulnerable to coextinction, and that maximum trophic level is a better predictor than other aspects of food web structure, such as species richness or trophic connectance. Extending this approach, we also find that maximum trophic level uncovers the relative vulnerability of ecosystem services to species coextinction across 12 empirical food webs.

## Introduction

Ecosystems are facing a wide variety of threats, including climate change, habitat loss, and invasive species, which can deteriorate ecosystems and disrupt ecological processes that support ecosystem services (Battisti et al., 2016). The direct risks of extinction from threats, such as land-use change and climate change, to select individual species and certain taxonomic groups are well-established (IPBES, 2019; Boyce et al., 2022; Penn & Deutsch, 2022; Thomas et al., 2004; Urban, 2015). While such threats are expected to continue leading to primary extinctions (Barnosky et al., 2011; Ceballos et al., 2020; Díaz et al., 2019), recent research points to the growing risk of coextinctions: additional secondary extinctions triggered by an initial primary extinction (Dunne & Williams, 2009; Kehoe et al., 2021; Morton et al., 2022; Strona & Bradshaw, 2022). For example, a specialist predator whose only prey species goes extinct due to a threat could also become extinct, or “coextinct”, even if the original threat does not directly impact the predator. Therefore, comprehending the full effect of ecosystem threats requires accounting for the potential coextinctions that may occur following primary extinctions due to lost interactions (Dunne & Williams, 2009; Koh et al., 2004), and the cumulative risk that both pose for ecosystem services (Keyes et al., 2021; Wilkes et al., 2024).

While recent studies predict that the risk of coextinctions *can* outweigh that of primary extinctions under climate and land-use change (Kehoe et al., 2021; Strona & Bradshaw, 2022), significantly less is known about the *circumstances or ecosystem characteristics under which* coextinction risk is expected to outweigh primary extinction risk across ecosystems more generally. Understanding when, or why, ecosystems may be especially prone to coextinctions is critical for accurately anticipating the extent of total species extinctions in the face of various threats. In food web ecology, the structure of trophic interactions between species within a food web has been studied extensively as a potential predictor of community vulnerability to coextinction. The number of species within a food web (size) and the density of interaction links between species (connectance) are two commonly studied aspects of food web structure, with conflicting evidence whether either structural metric promotes or hinders the occurrence of coextinctions (Dunne et al., 2002; Dunne et al., 2004; Keyes et al., 2024; Srinivasan et al., 2007; Staniczenko et al., 2010; Thébault et al., 2007). Multiple studies have also identified species at higher trophic levels as more vulnerable to extinction due to their dependence on species at lower trophic levels (Binzer et al., 2011; Häussler et al., 2020; Petchey et al., 2004), though it is less obvious whether vertical diversity, or the number of trophic levels in a food web, can predict aggregate coextinction risk for the entire community. Clarifying the relationships among structural features of food webs and coextinction risk could help identify ecosystems that are more susceptible to coextinctions.

Several methodological challenges impede a more nuanced understanding of which ecosystems are more likely to experience significant coextinction events. Widescale projections of coextinction rarely explicitly model the indirect effects of primary extinction based on the structure of known species interactions, instead relying on inferred species dependencies extrapolated from species trait and co-occurrence data (Koh et al., 2004; Strona & Bradshaw, 2018; Strona & Bradshaw, 2022; Strona et al., 2021). When such interaction data are available (often in the form of food webs), ecologists have typically used either topological approaches (Dunne et al., 2002; Dunne et al., 2004; Keyes et al., 2024) or dynamical models such as the allometric trophic network model (ATN; Yodzis & Innes, 1992) to predict outcomes from threats such as climate change (Eloranta et al., 2023; Gauzens et al., 2020), invasive species (Romanuk et al., 2009; Van Kleunen et al., 2023), and anthropogenic disturbance (e.g., fishing; Uusi-Heikkilä et al., 2021). Both have their disadvantages: topological models make the simplifying assumption that consumers are not at risk of coextinction until they have lost all their resources, thus underpredicting the extent of coextinction (Berg et al., 2015; Curtsdotter et al., 2011; Eklöf & Ebenman, 2006), while dynamical models require many parameters that are often difficult to obtain empirically, making them especially challenging to parameterize and validate for empirical ecosystems (Eklöf et al., 2013; Häussler et al., 2020).

There is a third approach, based on Bayesian networks, that is intermediate in complexity between topological and fully parameterized dynamical models. It can be thought of as a probabilistic generalization of the topological approach, relaxing modeling assumptions and expressing extinctions outcomes via probabilities instead of binary states (Figure 1). A critical improvement upon the topological approach is the ability to explicitly define consumers’ sensitivity to the *partial* loss of resources. Additionally, Eklöf et al. (2013) demonstrated that the Bayesian network approach can closely recreate results from the dynamical ATN model while requiring far fewer parameters. By balancing the realism of dynamical models with the simplicity of the topological approach, Bayesian networks can provide a useful baseline for modeling predictions of coextinction in food webs.

**Figure 1.**
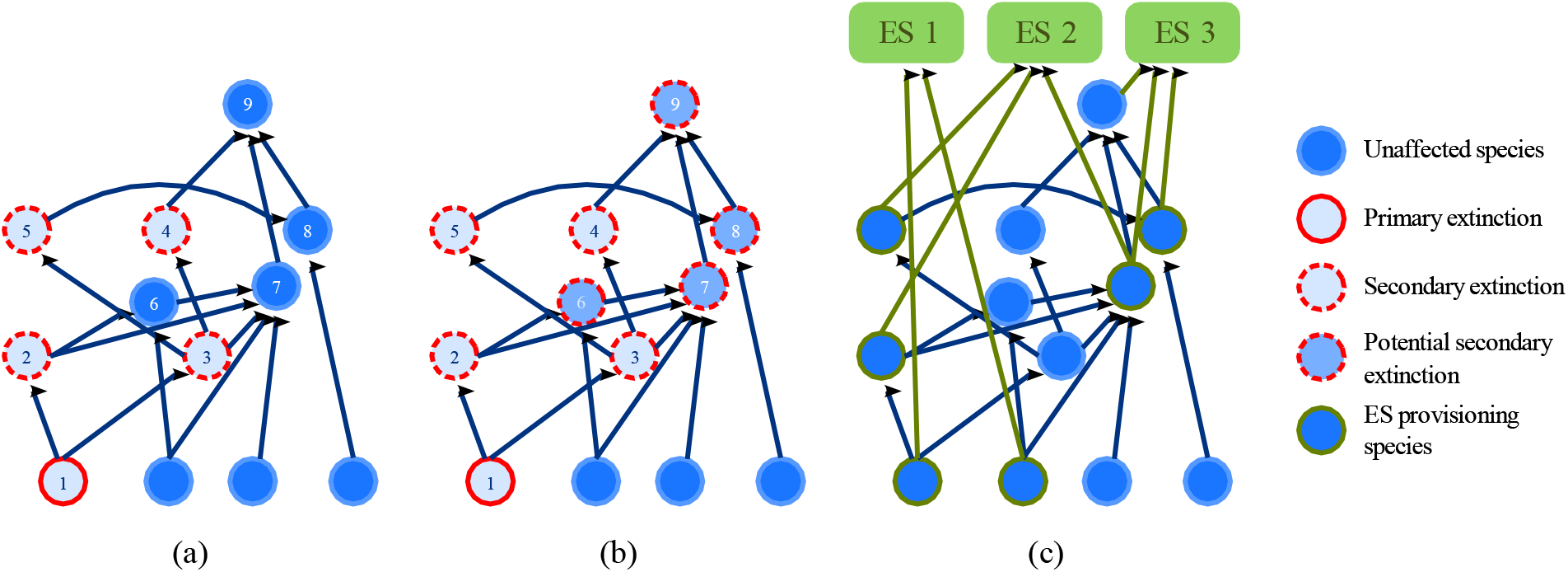
Comparison of extinction scenario simulation approaches (a and b), and food web augmented with ecosystem service (ES) nodes and links (c). In the topological approach (a), species 1 is chosen to first go extinct (solid red outline). Species 2 and 3 subsequently become extinct (dotted red outline) because they depend solely on species 1 as a resource. Species 4 and 5 also become subsequently extinct because they depend solely on species 3 as a resource. Importantly, the other species (species 6-9) are unaffected because they have at least one other available resource. In the Bayesian network approach (b), species 2-5 also go secondarily extinct following the extinction of species 1. However, the extinction probabilities of species 6-9 increase due to the partial loss of resource species. In the augmented food web (c), ES nodes are added to the food web (green nodes), and ES provisioning links (green links) connect ES providing species (solid green outline) to each ES.

Here, we ask which aspects of food web structure influence the importance of coextinction relative direct extinction risk, and how the vulnerability of communities to coextinction are subsequently passed on to the persistence of ecosystem services. To answer our questions, we present and demonstrate how the Bayesian network modeling approach can be used to quantify the importance of coextinction risk relative to direct extinction risk, and how it varies as a function of **1)** direct extinction risk; **2)** food web size and connectance; **3)** maximum trophic level of the food web (vertical diversity); and **4)** the degree to which consumers are sensitive to the loss of their resources.

We use a combination of mathematical analyses – deriving analytical solutions on simplified model food webs – and simulations on more complex model and empirical food webs. By comparing simulation outcomes for both model and empirical food webs, we evaluate whether synthetic food webs can be used to predict coextinction risk for real-world ecosystems. Additionally, we leverage recent advances that extended Bayesian networks to study the risk of coextinctions to the ecosystem services that depend on individual species embedded within a food web (Eklöf et al., 2025). Specifically, we quantify the risk for species coextinctions to negatively impact ecosystem services in 12 empirical food webs, examining a range of ecosystem service types (including provisioning, regulating, and recreational services) provided by species at various trophic levels. Identifying the characteristics of communities and threats that indicate high relative importance of coextinction could help guide conservation strategies, as managers could prioritize investment in acquiring the necessary data for ecosystems where coextinctions are expected to substantially outpace primary extinctions.

## Methods

### Quantifying the relative importance of coextinction

We simulated a range of extinction scenarios on food webs using the Bayesian network approach (Figure 1b; Jensen, 1996; Eklöf et al., 2013). To describe species’ baseline probabilities of extinction *ε_i_* due to a threat, we considered three threat profiles: a deterministic threat that affects all species equally (*ε_i_* = *ε* for all species *i*), a stochastic threat that affects each species differently (*ε_i_* ∼ Uniform[*ε* – 0.05, *ε* + 0.05] for all species *i*), and a deterministic basal-only threat that only affects basal species (*ε_i_* = *ε* for all basal species *i* and 0 for all non-basal species). For each threat profile, we simulated an extinction scenario for each value of *ε* varying from 0.1 to 0.9 in increments of 0.1.

The outputs of an extinction scenario simulation are the marginal extinction probabilities *p_i_* for each species *i* in the food web. By summing *ε_i_* over all species, we define the **Direct Effect** (*D*) of the threat as the expected number of primary extinctions that occur as a direct result of the threat, absent any coextinctions. Similarly, summing *p_i_* over all species yields the **Total Effect** (*T*) of the threat as the expected number of extinctions that occur either as a direct result of the threat or from subsequent coextinction (Table 1).

**Table 1.** Description of calculated quantities. Direct Effect and Total Effect describe the expected number of extinctions in a food web for a given extinction scenario simulation when coextinctions are not and are considered, respectively. Excess Extinction and Fragility are the two outcomes of interest in our analysis, describing different aspects of the relative importance of coextinction. Excess Loss of ES *s* describes the relative importance of coextinction of providing species for the continued provisioning of ecosystem service *s*.

| Quantity | Equation | Interpretation |
| --- | --- | --- |
| Direct Effect | $D = \sum_i \varepsilon_i$ | Expected number of extinctions to occur when effects of coextinction do not propagate throughout the food web |
| Total Effect | $T = \sum_i p_i$ | Expected number of extinctions to occur when the effects of coextinction propagate throughout the food web |
| Excess Extinction | $E = \frac{T}{D} = \frac{\sum_i p_i}{\sum_i \varepsilon_i}$ | Extent of coextinctions relative to primary extinctions, induced by the interaction structure of the food web |
| Fragility | $F = \frac{T - D}{S - D} = \frac{\sum_i p_i - \sum_i \varepsilon_i}{S - \sum_i \varepsilon_i}$ | Extent of food web collapse caused by coextinctions, induced by the interaction structure of the food web |
| Excess Loss of ES $s$ | $E_s = \frac{\sum_{i \in S} p_i}{\sum_{i \in S} \varepsilon_i}$ | Extent of coextinctions relative to primary extinctions, restricted to the set of species, $S$, that directly provide ecosystem service $s$ |

Using *D* and *T*, we define two outcome variables that quantify the relative importance of coextinction. The first outcome of interest, **Excess Extinction** (*E* = *T* / *D*), describes the number of coextinctions relative to primary extinctions. A high value of *E* indicates that the expected number of coextinctions is large relative to the expected number of direct extinctions – and thus corresponds to a greater relative importance of coextinction in the food web. The second outcome of interest, **Fragility** (*F* = (*T* – *D*) / (*S* – *D*), where *S* is the total number of species), quantifies the extent of food web collapse caused by coextinction. *F* always lies between 0 and 1; a high value of *F* indicates that coextinction is expected to remove a large portion of the remaining species that survive the initial wave of primary extinctions. To describe the relative importance of coextinction to the provisioning of an ecosystem service *s*, we define an analogous outcome of interest, **Excess Loss of Ecosystem Service *s*** (*E_s_*; Table 1).

### Simulating extinction scenarios

The marginal probability of extinction *p_i_* for each species *i* is determined by the form of a function, *w*(*f*), which describes consumers’ sensitivity to a loss of a fraction *f* of its resource species. For each consumer species *i*, its marginal probability of extinction *p_i_*(*f*) is calculated as

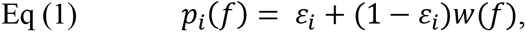

where *f* is the fraction of its resource species that have gone extinct. Because basal species do not require prey, their marginal probabilities of extinction are *p_i_* = *ε_i_*. In Equation 1, *w*(*f*) is a monotonically increasing function from 0 at *f* = 0 to 1 at *f* = 1. We choose *w*(*f*) to be one of three different functional forms (Figure 2): *w*(*f*) = *f* (linear), *w*(*f*) = *f*^2^ (convex), and *w*(*f*) = 1 – (1 – *f*)^2^ (concave). Consumers are less sensitive to a partial loss of resources when *w*(*f*) is convex and are more sensitive when *w*(*f*) is concave.

**Figure 2.**
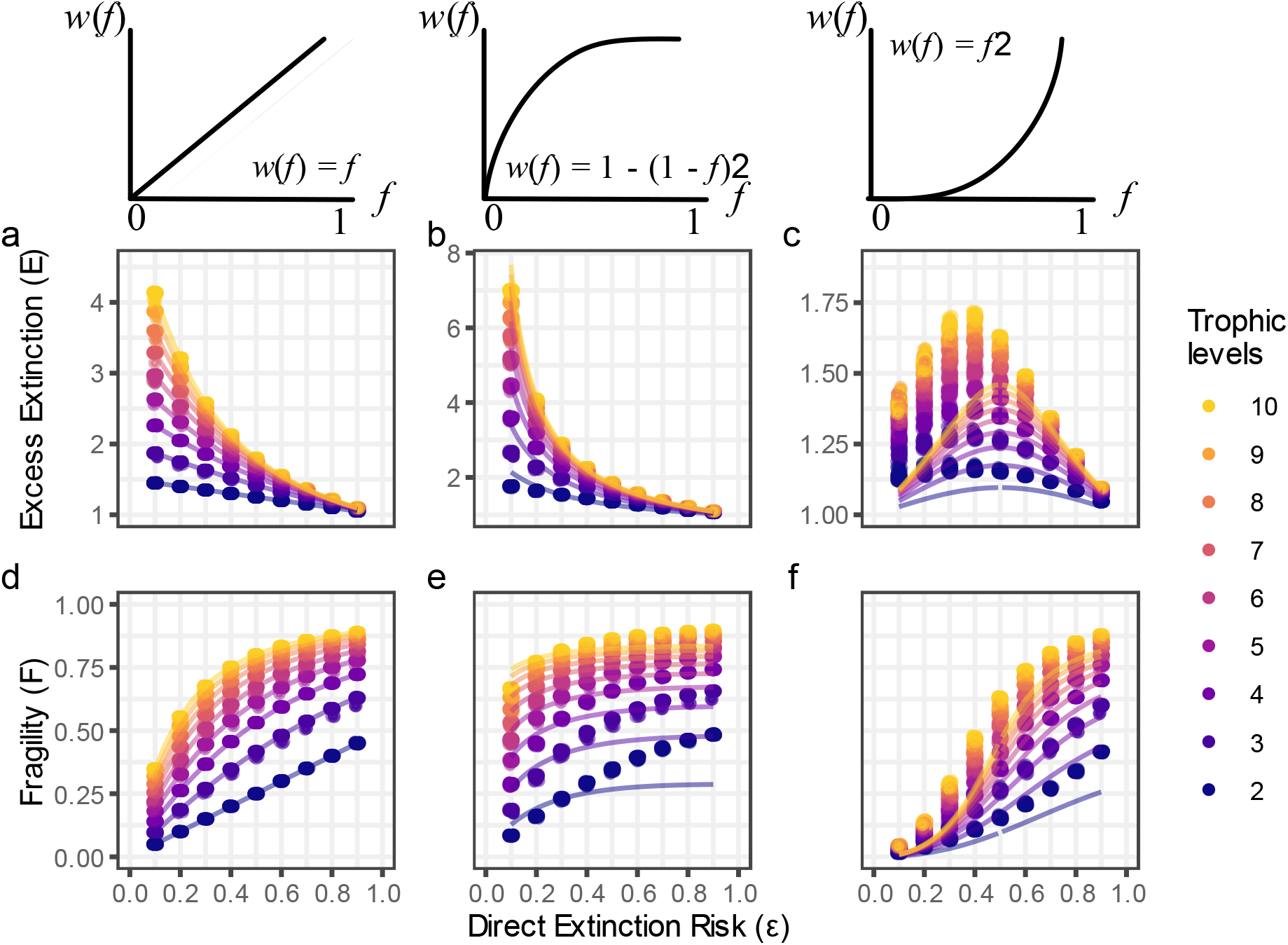
Excess Extinction and Fragility as a function of direct extinction risk in stratified food webs. Excess Extinction (top row) and Fragility (bottom row) as a function of direct extinction risk (ε) with a deterministic threat. Each point describes the outcome of an extinction simulation (horizontal jitter added to reduce overplotting) when the response function *w*(*f*) is linear (left column), concave (middle column), and convex (right column). For the linear response function case, each line graphs Excess Extinction and Fragility as calculated by Equations 2 and 3, respectively. For the concave and convex cases, each lines graphs approximations of Excess Extinction and Fragility as calculated by equations in Supporting Information. Point color indicates *m*, the number of trophic levels in the stratified food web.

### Food webs

We conducted extinction scenario simulations on a suite of food webs along of spectrum of complexity: stratified (“rectangular”) food webs (Figure S1a; Borrvall et al., 2000), niche model food webs (Figure S1b; Williams & Martinez, 2000), and empirical food webs.

We first generated stratified food webs, in which *n* species are assigned to each of *m* discrete trophic levels, for a total species richness of *S* = *m*·*n*. Feeding interactions are restricted to only occur between species in adjacent trophic levels, precluding any omnivory. We also assumed that species within trophic levels are interchangeable and that consumers have no preference among their resources. This assumption is expressed by setting a single interaction probability *q* between species in adjacent trophic levels. We simulated stratified food webs with *m* varying from 2 to 10 trophic levels, while fixing *n* = 10 species per trophic level and interaction probability *q* = 0.5. This led to overall food web connectance values between 0.0045 and 0.013. We generated 50 replicates for each combination of parameters, resulting in a total of 450 stratified food webs.

We next generated niche model food webs (Williams & Martinez, 2000), which allow for omnivory. Niche model food webs have been used extensively in the study of food web structure and stability, and have been shown to reliably recreate multiple aspects of empirical food web structure (Cirtwill & Wootton, 2022; Domínguez-García et al., 2019; Dunne & Williams, 2009; Johnson et al., 2014; Stouffer & Bascompte, 2010; Williams & Martinez, 2000). We simulated niche model food webs with size *S* = 25, 50, 100, 150, and 200 species, and connectance *C* = 0.05, 0.10, 0.15, and 0.20. Due to the stochastic nature of niche model food web generation, these target connectance values were achieved with a 5% error tolerance. We again generated 50 replicates for each combination of parameters, resulting in a total of 1,000 niche model food webs.

Additionally, we examined 12 well-studied empirical food webs that span multiple ecosystem types (Table S1; Baird & Ulanowicz, 1989; Goldwasser & Roughgarden, 1993; Hechinger et al., 2011; Martinez, 1991; Mendonça et al., 2018). As with our synthetic food webs, these empirical food webs varied in their size (31 to 105 species) and connectance (0.072 to 0.175). To evaluate whether the niche model could accurately predict extinction scenario simulation outcomes for empirical food webs, we generated 50 niche model food webs parameterized by the same size and target connectance as each of the 12 empirical food webs. Using available data linking ecosystem services (ES) to species directly providing ecosystem services (Dee & Keyes, 2022; Keyes, 2023; Keyes et al., 2021; Wilkes et al., 2024), we supplemented each empirical food web with additional ecosystem service nodes and links (Figure 1c).

We calculated Excess Extinction (*E*) and Fragility (*F*) for each extinction scenario simulation, comprised of a food web, a threat profile, a baseline probability of extinction ε, and a functional form of *w*(*f*). For empirical food webs, we additionally calculate Excess Loss of ES *s* (*E_s_*) for all documented ecosystem services *s*.

### Applicability of analytical solutions, from simple to complex food webs

The simplified structure of stratified food webs allowed us to find closed-form solutions for Excess Extinction *E* and Fragility *F* when consumers have a linear response function *w*(*f*) = *f* and face a deterministic threat. Given the direct extinction risk ε and the number of trophic levels *m* in the food web, we get (Supporting Information):

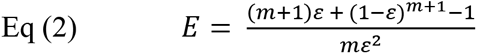

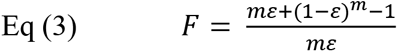

The two are linked by

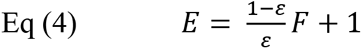

Furthermore, we derived closed-form solutions for approximations of *E* and *F* when consumers have a non-linear response function *w*(*f*), though this required imposing an additional assumption on the number of species per trophic level (Supporting Information). Though we solved for *E* and *F* under the assumption of a deterministic threat, we find that our solutions closely approximate the behavior of *E* and *F* under a stochastic threat as well (Figures S10-S15).

Our analytical solutions indicate that when there is a deterministic threat and a linear response function, simulation outcomes for stratified food webs should be predicted by just the direct extinction risk and the number of trophic levels in the food web. We test whether the vertical diversity of a food web is also a relevant predictor of coextinction outcomes for more complex food webs. To do so, we calculated the maximum prey-averaged trophic level (Levine, 1980) for each niche model food web and empirical food web. We then estimated the parameter *m\** that best fit the observed values of Excess Extinction *E* obtained from extinction scenario simulation outcomes, constrained to the functional form of Equation 2. We define *m\** as the food web’s effective trophic level, the equivalent number of trophic levels necessary in a stratified food web to yield the same values of *E* and *F* in extinction scenario simulations. By comparing a complex food web’s true maximum trophic level to its effective maximum trophic level (Figure 4 left panel), we examine whether vertical diversity can be a useful predictor of coextinction outcomes in complex food webs, despite significant differences in food web structure and complexity relative to that of stratified food webs.

## Results

### Excess Extinction and Fragility in stratified food webs

For a deterministic threat and a linear response function, we find that Excess Extinction *E* decreases with direct extinction risk *ε* but increases with the number of trophic levels *m* in the food web (Figure 2a). On the other hand, Fragility *F* increases with both direct extinction risk and the number of trophic levels (Figure 2d). Thus, coextinctions comprise a larger proportion of the total number of extinctions when direct extinction risk ε is small, but the absolute extent of coextinctions in the community is greater when ε is large. In food webs with many trophic levels (high vertical diversity), coextinctions outpace primary extinctions and there are also fewer species remaining after coextinctions have propagated through the community.

By contrast, *E* and *F* are independent of *q* (the probability of species interaction) in stratified food webs, as well as the number of species per trophic level *n* (Equations 2 and 3). Thus, for stratified food webs facing a deterministic threat with a linear response function, the relative importance of coextinction does not vary with food web connectance and varies with food web size only through changes in the number of trophic levels.

Direct extinction risk, vertical diversity, size, and connectance have the same qualitative effects on Excess Extinction and Fragility when consumers are more sensitive to a partial loss of resources (Figures 2b and 2e). However, when consumers are less sensitive to a partial loss of resources, Excess Extinction no longer decreases monotonically with direct extinction risk and is instead greatest at moderate values of direct extinction risk (Figure 2c). Thus, the effect of direct extinction risk on Excess Extinction can vary dramatically depending on how consumers respond to a partial loss of their resources.

### Excess Extinction and Fragility in niche model food webs

In niche model food webs, we find the exact same qualitative relationship between Excess Extinction *E* and direct extinction risk ε as in stratified food webs (Figure 3): under a deterministic threat and a linear response function, Excess Extinction decreases as direct extinction risk increases. Similarly, as with stratified food webs, Fragility also increases with direct extinction risk (Figures S2 and S3).

**Figure 3.**
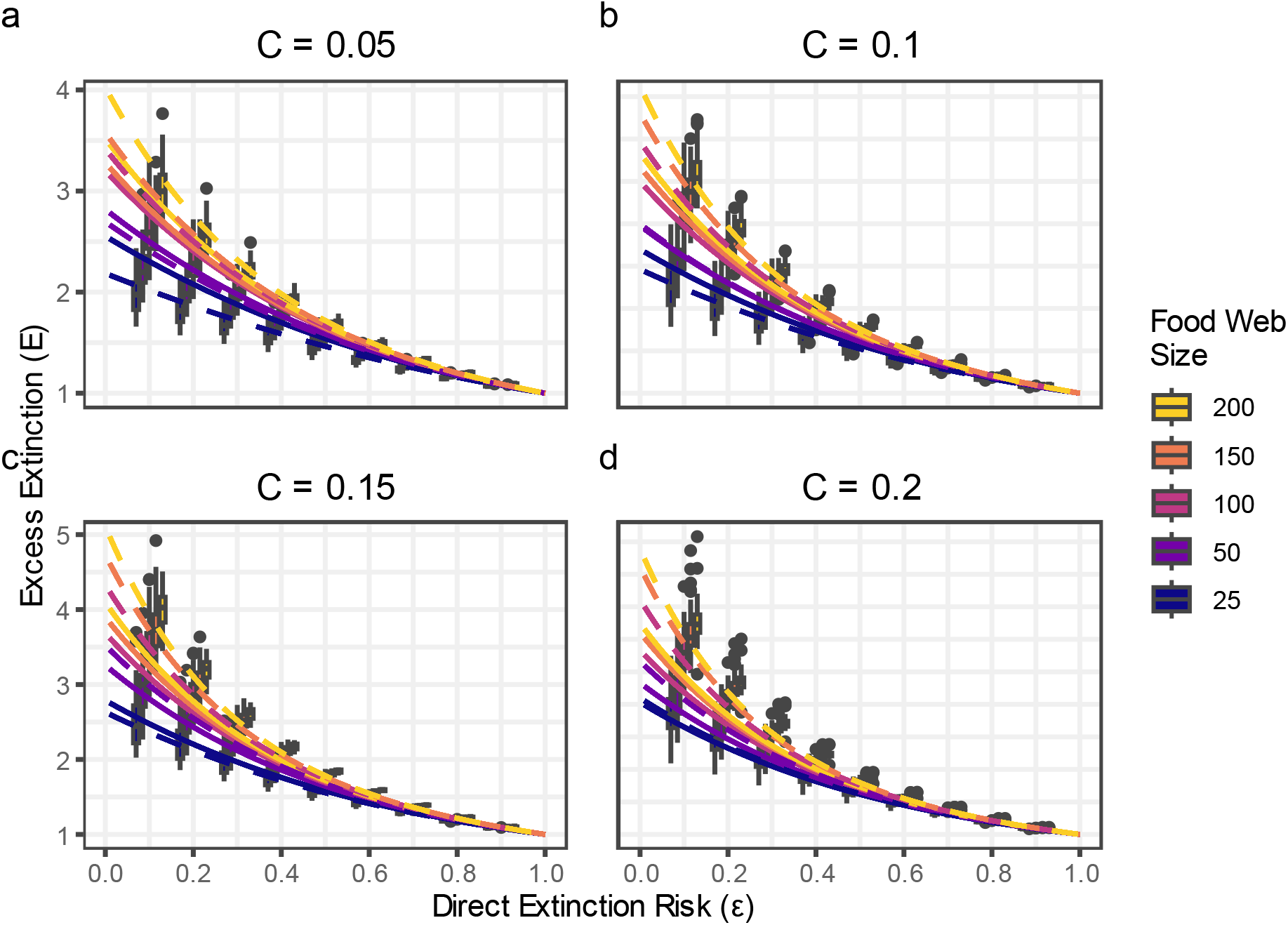
Excess Extinction as a function of direct extinction risk in niche model food webs with different connectance values. Excess Extinction as a function of direct extinction risk (ε) with a deterministic threat and linear response function *w*(*f*). Plots are faceted by food web connectance, and boxplot color indicates food web size. Lines graph Excess Extinction as calculated by Equation 2, parameterized by true maximum trophic level (solid) and the effective maximum trophic level (dashed) estimated by the line of best fit (see Figure 4 left panel).

**Figure 4.**
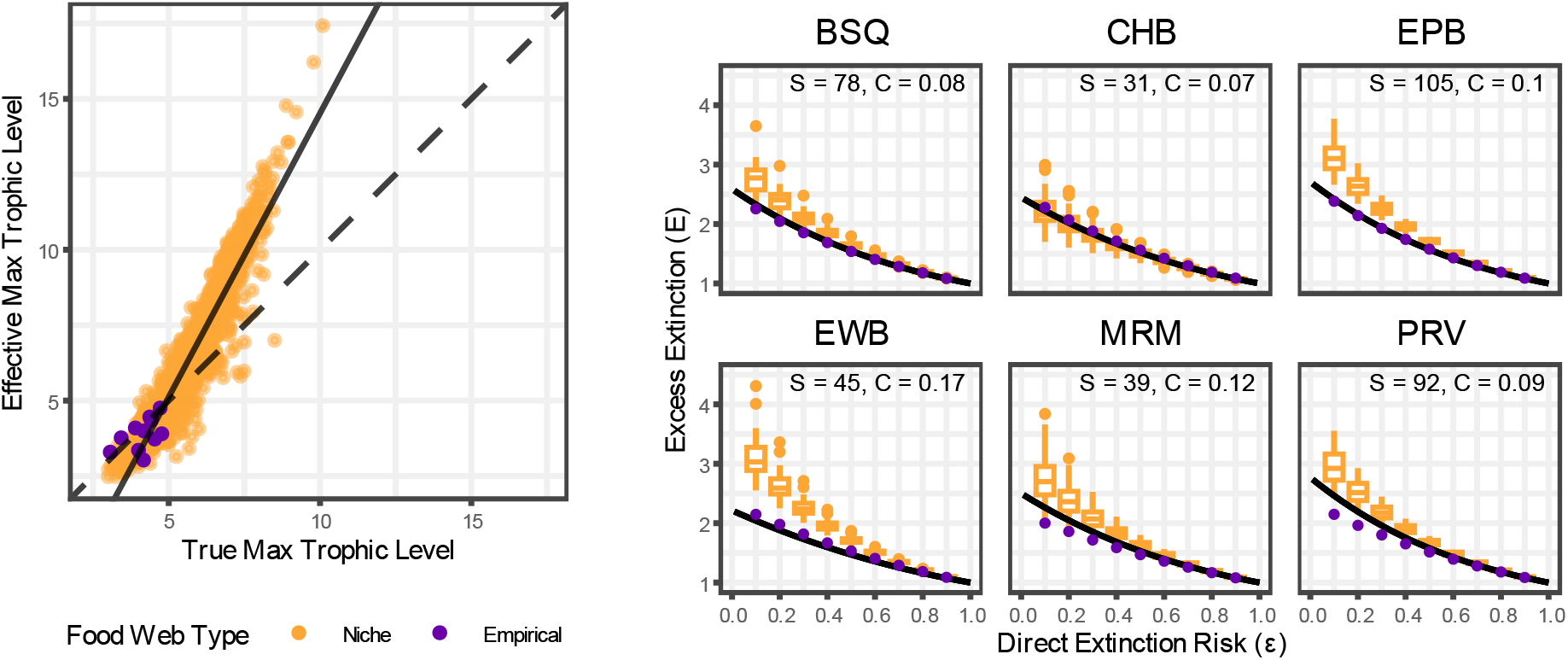
Comparison of true maximum trophic level and effective maximum trophic level in niche model and empirical food webs, and extinction simulation outcomes for empirical food webs compared to corresponding model food webs. For each extinction simulation on niche model and empirical food webs with linear response function *w*(*f*) and a deterministic threat, we calculated the maximum trophic level of the food web (left panel abscissa) as well as the parameter *m\** that best fits the observed value of *E* using Equation 2 (left panel ordinate). The solid line is the line of best fit. The dotted line is *y* = *x*, representing the points where the true maximum trophic level of a food web coincides with its effective maximum trophic level.Extinction simulation outcomes using a linear response function *w*(*f*) and a deterministic threat are shown for a subset of empirical food webs and their corresponding model food webs. Purple points indicate *E* for outcomes on empirical food webs. Golden box plots indicate the range of simulation outcomes for corresponding niche model food webs parameterized by the size and connectance of each empirical food web. Black lines graph *E* as calculated by Equation 2, parameterized by the maximum trophic level of each empirical food web. See Figure S16 for plots for all empirical food webs.

Excess Extinction and Fragility also increase with both food web size and connectance, although these dependencies are relatively weak (Figures S2 and S3). The number of trophic levels in niche model food webs increases with both food web size and connectance (Figure S7); thus both Excess Extinction and Fragility increase with the number of trophic levels in niche model food webs (Figures S12 and S13).

Direct extinction risk, food web size, and food web connectance have the same qualitative relationship with Excess Extinction and Fragility in the linear and concave cases (Figures S12 and S13), i.e. when species are more sensitive to a partial loss of resources. However, when species are less sensitive to a partial loss of resources (the convex case) and direct extinction risk is low, Excess Extinction tends to decrease with food web size (Figure S2c).

### Excess Extinction and Fragility in empirical food webs

Mirroring previous findings in synthetic food webs, Excess Extinction *E* decreases with direct extinction risk *ε* (Figure S14a) while Fragility *F* increases with direct extinction risk *ε* (Figure S15a) under a deterministic threat and a linear response function. As with stratified webs, food web size and connectance do not have clear effects on *E* and *F*, with *E* and *F* remaining relatively constant for a given direct extinction risk across the entire range of food web size (Figures S4a and S4e, respectively) and connectance (Figures S4c and S4g, respectively).

The dependence of *E* and *F* on direct extinction risk is similar when consumers are less sensitive to a partial loss of resources, but in this case, we observe that Excess Extinction tends to decrease with both food web size and connectance (Figures S4b and S4d, respectively).

### Predicting the relative importance of coextinction in complex food webs using stratified webs

We observe that the relationship between food webs’ true and effective maximum trophic level (Figure 4 left panel) is remarkably linear and tightly coupled (*ρ* = 0.934). This relationship between maximum and effective trophic level is particularly well-approximated for realistic ranges of trophic levels (i.e., between 3.07 and 4.77 in our empirical food webs). While true trophic level and effective trophic level covary predictably, the two do not coincide exactly (Figure 4 left panel, dashed line). We find that effective trophic level is less than true trophic level when the maximum trophic level of the complex food web is low. In this case (generally the case for small food webs), using the true maximum trophic level of a food web to predict extinction outcomes will likely lead to overestimations of *E* and *F*. Conversely, when the maximum trophic level of a food web is high (generally the case for large food webs), effective trophic level is greater than true trophic level and thus predictions made by the true maximum trophic level will generally underestimate *E* and *F*.

Comparing our analytical solutions to outcomes generated by each empirical web’s corresponding niche model food webs, we observe that Excess Extinction for empirical food webs is better predicted by stratified food webs parameterized by only the true maximum trophic level of the empirical food web (Figure 4 right panel) than by niche model food webs parameterized by the size and connectance of the web: in almost all cases, Excess Extinction is lower in empirical food webs than predicted by corresponding niche model food webs (Figure 4 right panel, Figure S16). However, four rocky intertidal food webs from Mendonça et al. (2018) were predicted poorly by corresponding niche model food webs, and not particularly well (though still much better than niche webs) by corresponding stratified webs: Porto da Cruz, Portugal; Canico, Portugal; Cabo Raso, Portugal; and Raio Verde, Portugal (Figure 4 MRM and PRV; Figure S16 MPC, MRM, PCR, and PRV). We did not find that variation in the extent of omnivorous interactions could sufficiently explain differences in the predictive performance of corresponding stratified food webs for empirical food webs (see Supporting Information and Figure S8).

### Modeling the loss of ecosystem services

We quantified the relative importance of species coextinction for the persistence of eight ES across 12 empirical food webs (Figure 5). Under a deterministic threat and a linear response function, we found that the Excess Loss of ES *s* (*E_s_*) decreases with the direct extinction risk to species in the food web. Furthermore, within a given food web, *E_s_* tends to be greater for services provided by species at higher trophic levels (such as fishing and birdwatching) than for services provided by basal species (such as carbon sequestration and wave attenuation). However, there is variation in the absolute value of *E_s_* for a given ecosystem service across food webs. For example, the relative importance of species coextinction is greater for the persistence of recreational fishing in the Wembury, England food web (Figure 5 EWB) than in the Porto da Cruz, Portugal food web (Figure 5 MPC).

**Figure 5.**
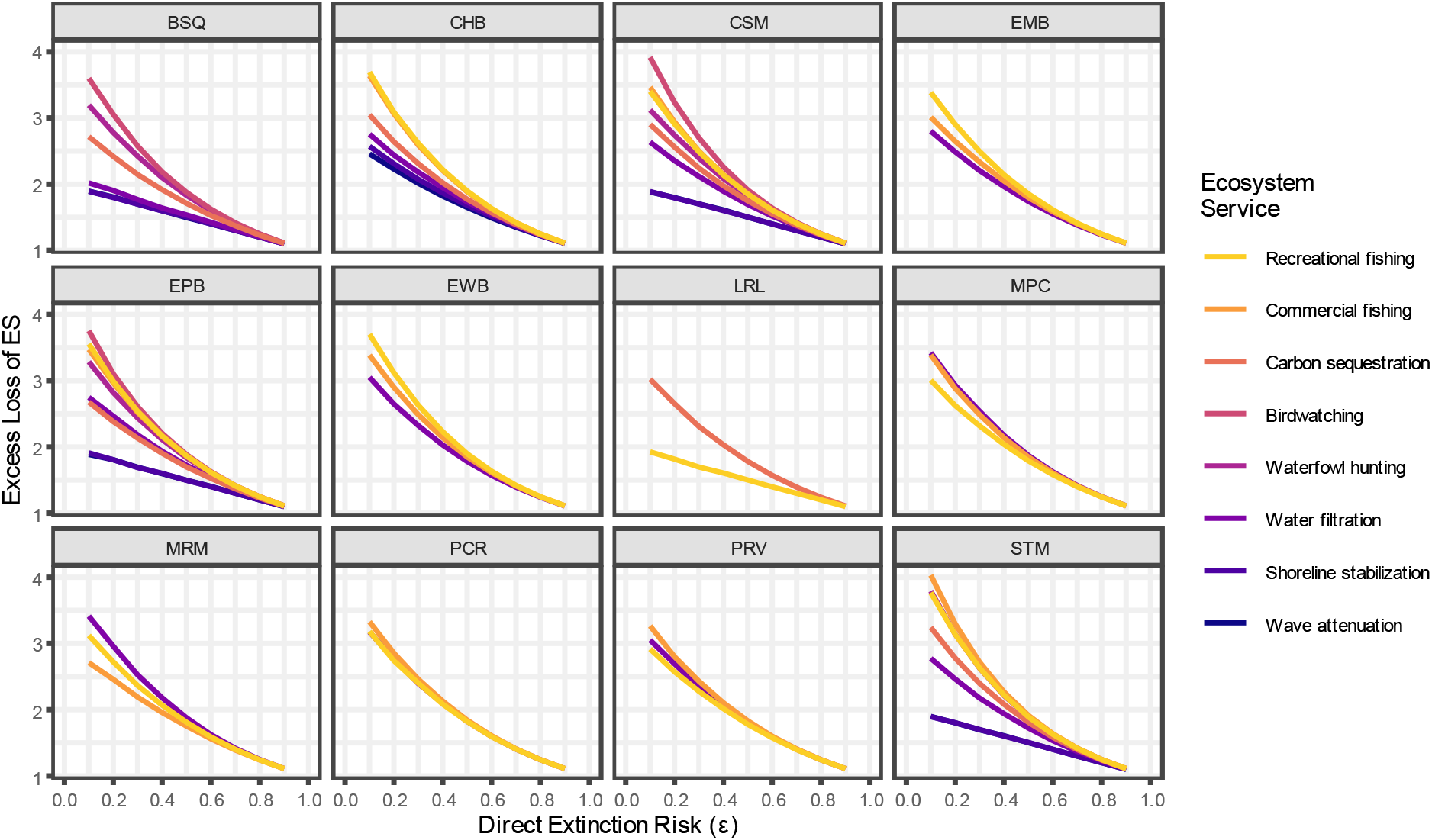
Excess Loss of Ecosystem Services, grouped by empirical food web. Simulation outcomes for Excess Loss of ES are plotted as a function of direct extinction risk, under a deterministic threat and a linear response function *w*(*f*). Some empirical food webs documented ecosystem service provisioning data more comprehensively than others, leading to variation in the number of ecosystem service loss simulations performed per empirical food web.

## Discussion

Our study leverages an underused modeling framework, Bayesian networks, to identify structural characteristics of food webs that are conducive to cascading coextinctions. Combining studies investigating the increasing importance of coextinctions (Kehoe et al., 2021; Strona & Bradshaw, 2018; Strona & Bradshaw, 2022) with recent advances for modeling food web dynamics (Eklöf et al., 2013; Eklöf et al., 2025), we aim to identify factors that can indicate whether a community is especially vulnerable to coextinctions. Through our analysis of simple stratified food webs, we identified direct extinction risk, the vertical diversity of the food web, and consumers’ response to partial resource loss as the most salient factors in determining the importance of coextinctions relative to primary extinctions. Notably, these factors remained the most important predictors for niche model and empirical food webs, despite substantial differences in the complexity of food web structure relative to stratified food webs.

Though Bayesian networks require fewer parameters than dynamic models, it is nevertheless a challenge to construct accurate and comprehensive empirical food webs. For this reason, we explored the extent to which simplified food webs and proxies of food web structure (such as maximum trophic level) could sufficiently predict extinction simulation outcomes in complex food webs. A key insight from our analytical results (and confirmed by simulations) is that vertical diversity – the maximum number of trophic levels in a food web – is the key indicator of community vulnerability to coextinction, as measured by Excess Extinction and Fragility. Previous work has found that communities containing longer food chains (i.e., greater vertical diversity) were less stable, as measured by length of recovery time (Pimm & Lawton, 1977; Zhao et al., 2019) and by the leading eigenvalue of the Jacobian matrix (Vagnon et al., 2023). Thus, it seems that communities with greater vertical diversity may be considered unstable along multiple dimensions. Importantly, the consumers’ trophic positions (and thus the vertical diversity of a community) can be determined empirically through stable isotope techniques (Post, 2002), even when the exact structure of trophic interactions is unknown.

We found that knowing the maximum trophic level of an empirical food web was usually sufficient to predict coextinction outcomes well, but this was not always true. For four of the 12 empirical food webs (Porto da Cruz, Portugal; Canico, Portugal; Cabo Raso, Portugal; Raio Verde, Portugal), relying on the maximum trophic level to predict coextinction outcomes led to overestimates of coextinction. We did not find any systematic differences in food web structure between these four food webs and the other eight, in terms of size, connectance, and omnivory. Future investigations could explore more sophisticated measures of food web structure to establish whether there are structural features of food webs that indicate how closely coextinction outcomes are expected to match predictions based on maximum trophic level.

Recent findings documenting the growing risk of coextinction generally rely on trait- and location-based data as proxies for inferring species dependencies (Strona & Bradshaw, 2018; Strona & Bradshaw, 2022; Strona et al., 2021). Studies that do utilize food webs frequently use a topological approach to model coextinctions (Dunne et al., 2002; Dunne et al., 2004; Keyes et al., 2024), but neither of these approaches can account for consumer sensitivity to the partial loss of their resources. Our study shows that increased consumer sensitivity leads to increases in the total portion of species lost to coextinction (Fragility), as well as increases in the number of coextinctions relative to primary extinctions (Excess Extinction). Since the topological approach represents the case in which consumers are completely insensitive to a partial loss of resources (Eklöf et al., 2013), it is likely that topological models underestimate the extent of coextinctions that occur. While the empirical data needed to describe this consumer sensitivity can be unavailable or difficult to collect, our results suggest that estimating this characterization is imperative for accurately predicting the extent of coextinctions within a community.

Recent work has shown that food web robustness to coextinction is highly correlated to ES robustness, yet this association is likely context-dependent (Keyes et al., 2021). We found that the Excess Loss of ecosystem services tends to be greater for services provided by species at higher trophic levels (consistent with Jacob et al., 2011) and are thus more likely to be lost due to species coextinctions. This mirrors our findings that species at higher trophic levels are at greater risk of coextinction. Furthermore, the same ecosystem service may have differing risks of being lost in different food webs, highlighting the fact that system-specific knowledge about both species interactions and service provisioning is necessary to accurately assess the risk of losing a given ecosystem service. Analogous to our investigation of the functional form of consumers’ response to a partial loss of resources, future work could explore the functional form of services’ response to a partial loss of provisioning species and potential associations between ecosystem service type and functional form.

The Bayesian network approach has several limitations. First, parameterizing Bayesian networks may still be difficult for real-world threats and communities. For instance, data needed to quantify the direct extinction risk for each species from a particular threat is often unavailable. One possible solution is to estimate direct extinction risks using empirical population-level data such as mean population size and variability of population size over time (O’Grady et al., 2004), or to use trait-based proxies such as habitat range size, body size, and fecundity (Chichorro et al., 2019; González-Suárez et al., 2013; Murray et al., 2010).

Another drawback is the exclusion of top-down effects. Global change, specifically warming, can amplify top-down control exerted by predators (Barton et al., 2009; Shurin et al., 2012; Vidussi et al., 2011). As ecosystems experience warming and other environmental changes in the future, accounting for top-down effects could become increasingly important. Because the strength of top-down effects can be highly dependent on disturbance characteristics and community composition (Baum & Worm, 2009), as well as highly variable over time (Fox, 2007), it is difficult to predict the extent to which this omission could alter our findings.

In addition, our investigation of food web vulnerability to coextinction did not account for interaction rewiring – that is, predators were unable to trophically adapt by establishing new feeding links following the extinction of their prey. Ecosystems are predicted to experience more frequent interaction rewiring due to influxes of invading generalist species (Bartley et al., 2019; Korstch et al., 2015), since generalists are also those most likely to rewire their trophic interactions (Lázaro and Gómez-Martínez, 2022). While theoretical studies have found mixed effects of interaction rewiring on community vulnerability to coextinction (Gilljam et al., 2015; Staniczenko et al., 2010), the prevalence of rewiring and its effects on stability in real-world communities remain underexplored (but see Polazzo et al., 2022).

Fortunately, the Bayesian network approach can be extended to allow for rewiring (Eklöf et al., 2013). Since this increases the number of required parameters and makes mathematical analysis of even simplified food webs potentially intractable, future studies of Bayesian networks based on simulation could greatly advance our understanding of the role that rewiring plays in altering coextinction outcomes.

## Conclusions

As the concern of coextinction is expected to continuously increase under climate change and other anthropogenic drivers (Kehoe et al., 2021; Strona & Bradshaw, 2018; Strona & Bradshaw, 2022), identifying the ecosystems and/or interactions most vulnerable to significant alteration will become increasingly critical. To that end, we explored an underused modeling framework, Bayesian networks, to identify conditions under which ecosystems with interacting species could be more vulnerable to coextinctions, relative to primary extinctions. Our findings suggest that critical predictors of the relative importance of coextinction include the number of trophic levels in a food web and the sensitivity of consumers to a partial loss of resources. The vertical diversity of an ecosystem can be an instrumental indicator of both the importance of coextinctions relative to primary extinctions (Excess Extinction) as well as the total portion of species that will be lost (Fragility). Our findings also highlight the fact that how consumers respond to partial resource loss can significantly impact the extent of coextinction and should be accounted for in any evaluation of coextinction risk within an ecosystem. Together, these results illustrate that Bayesian networks provide a valuable framework for modeling risks of species coextinction and ecosystem service provisioning losses, warranting further study as a fruitful alternative to the topological and dynamical models commonly used by researchers.

## Supporting information

SuppInfo_proofs

SuppInfo_text_figures_tables

## Authorship statement

HL, GB, and LED conceptualized the study and designed the methodology. HL and GB conducted the analysis. AE and LED contributed to refining study goals and methodology. HL prepared the initial draft of the manuscript, and all authors contributed substantially to revisions.

## Data accessibility statement

The data and code used to generate the findings of this study are openly available on Github at https://github.com/heli8222/BayesianNetworkCoextinction and archived in Zenodo at https://doi.org/10.5281/zenodo.21939117.

## Acknowledgements

L. Dee and H. Li acknowledge support from NSF Biological Oceanography award #2049360 and the Eppley Foundation.

## Notes

https://github.com/heli8222/BayesianNetworkCoextinction

https://doi.org/10.5281/zenodo.21939116

### Competing Interest Statement

The authors have declared no competing interest.

### Funding

- National Science Foundation

