## Supplementary material for "Maximum trophic level predicts food webs’ susceptibility to coextinctions": SuppInfo_proofs

### Supporting Information Proofs

Analysis of stratified food webs using Bayesian networks

#### 1 Extinction probability of a single consumer

We start from a simple network with $n$ resource species $R_1, ..., R_n$ and a single consumer, $C$, connected to all of them. The resource species are not necessarily basal. Let the direct extinction risk be the same $\epsilon$ for all species. We denote the marginal probability of extinction of species $i$ with $p_i$. Assuming that this marginal extinction probability is the same for all resources ($p_{R_1} = p_{R_2} = ... = p_{R_n} \equiv p_R$), what is the consumer's marginal probability of extinction, $p_C$?

We can express this as follows:

$$\begin{aligned}
p_C = {} & P(\neg C|R_1 R_2 \dots R_n)\,\underbrace{(1-p_{R_1})(1-p_{R_2})\cdots(1-p_{R_n})}_{(1-p_R)^n,\text{ since } p_{R_i}\equiv p_R} \\
& + nP(\neg C|\neg R_1 R_2 \dots R_n)\,\underbrace{p_{R_1}(1-p_{R_2})\cdots(1-p_{R_n})}_{p_R(1-p_R)^{n-1}} \\
& + \binom{n}{2} P(\neg C|\neg R_1 \neg R_2 \dots R_n)\,\underbrace{p_{R_1}p_{R_2}\cdots(1-p_{R_n})}_{p_R^2(1-p_R)^{n-2}} \\
& + \cdots \\
& + nP(\neg C|R_1 \neg R_2 \dots \neg R_n)\,\underbrace{(1-p_{R_1})p_{R_2}\cdots p_{R_n}}_{p_R^{n-1}(1-p_R)} \\
& + P(\neg C|\neg R_1 \neg R_2 \dots \neg R_n)\,\underbrace{p_{R_1}p_{R_2}\cdots p_{R_n}}_{p_R^n}.
\end{aligned} \tag{1}$$

In this setup, the resources are assumed equivalent from the point of view of the consumer. Therefore the conditional probabilities $P(\neg C|R_1 R_2 \dots R_n)$ etc. only depend on the fraction of absent resources. Using the notation $P(\neg C|i)$ for the probability that $C$ is extinct given that $i$ of its resource species are absent, we can rewrite Eqn 1 as

$$\begin{aligned}
p_C = {} & P(\neg C|0)p_R^0(1-p_R)^n \\
& + nP(\neg C|1)p_R^1(1-p_R)^{n-1} \\
& + \binom{n}{2} P(\neg C|2)p_R^2(1-p_R)^{n-2} \\
& + \cdots \\
& + nP(\neg C|n-1)p_R^{n-1}(1-p_R)^1 \\
& + P(\neg C|n)p_R^n(1-p_R)^0,
\end{aligned} \tag{2}$$

or

$$p_C = \sum_{i=0}^{n} \binom{n}{i} P(\neg C|i) p_R^i (1-p_R)^{n-i} \tag{3}$$

more compactly.

The conditional probabilities $P(\neg C|i)$ are expressed via

$$P(\neg C|i) = \epsilon + (1-\epsilon) w\left(\frac{i}{n}\right), \tag{4}$$

where $i/n$ is the fraction of resource species that are absent, and $w$ is some monotonically increasing function such that $w(0) = 0$ and $w(1) = 1$. Substituting this into Eqn 3, we get

$$\begin{aligned} p_C &= \sum_{i=0}^{n} \binom{n}{i} \left[\epsilon + (1-\epsilon) w\left(\frac{i}{n}\right)\right] p_R^i (1-p_R)^{n-i} \\ &= \epsilon \sum_{i=0}^{n} \binom{n}{i} p_R^i (1-p_R)^{n-i} + (1-\epsilon) \sum_{i=0}^{n} w\left(\frac{i}{n}\right) \binom{n}{i} p_R^i (1-p_R)^{n-i}. \end{aligned} \tag{5}$$

In the first term, the sum is over the binomial distribution and is therefore 1:

$$p_C = \epsilon + (1-\epsilon) \sum_{i=0}^{n} w\left(\frac{i}{n}\right) \binom{n}{i} p_R^i (1-p_R)^{n-i}. \tag{6}$$

To make further progress, we must specify the function $w$. We do so below for three choices:

- Linear case, with $w(i/n) = i/n$ (Section 2);
- Convex nonlinear case, with $w(i/n) = (i/n)^2$ (Section 3);
- Concave nonlinear case, with $w(i/n) = 1 - (1 - (i/n))^2$ (Section 4).

#### 2 Excess Extinction and Fragility in the linear case

With the choice $w(i/n) = i/n$, Eqn 6 reads

$$p_C = \epsilon + (1-\epsilon) \sum_{i=0}^{n} \frac{i}{n} \binom{n}{i} p_R^i (1-p_R)^{n-i}. \tag{7}$$

Bringing the constant factor $1/n$ in front of the summation:

$$p_C = \epsilon + \frac{1-\epsilon}{n} \sum_{i=0}^{n} i \binom{n}{i} p_R^i (1-p_R)^{n-i}. \tag{8}$$

The sum is now simply the mean of the binomial distribution, $np_R$, so the expression simplifies to

$$p_C = \epsilon + (1-\epsilon) p_R. \tag{9}$$

This is independent of the number of connections $n$. As a consequence, any consumer species preying on any subset of the resources $R_1, \ldots, R_n$ will have its marginal extinction probability be given by Eqn 9.

Extrapolating this idea, let us imagine a stratified trophic chain. At each trophic level $k$ the species may only feed on species at level $k-1$ and are eaten only by those at level $k+1$. We specify that each level has $n$ species, although due to the independence of Eqn 9 from $n$, this assumption is easy to relax. Each species has a direct extinction risk of $\epsilon$. Then at level 1, each species has $\epsilon$ as their marginal extinction probability as well—so all species at level 1 have the same extinction probability, $p_1 = \epsilon$. Since these are all equal, it follows from Eqn 9 that all probabilities at level 2 are also equal: $p_2 = \epsilon + (1-\epsilon)p_1 = \epsilon + (1-\epsilon)\epsilon = 1-(1-\epsilon)^2$. And so on: the same idea applies recursively to all levels. This means that Eqn 9 can be generalized and written as a recursion giving the probabilities at the next level given those at the previous one:

$$p_{k+1} = \epsilon + (1-\epsilon)p_k, \tag{10}$$

where $p_k$ is the marginal extinction probability of any one species at trophic level $k$. The initial condition to this recursion is $p_1 = \epsilon$.

This is a linear recurrence equation that is easy to solve explicitly:

$$p_k = 1-(1-\epsilon)^k. \tag{11}$$

Indeed, for $k = 1$ this yields $p_1 = \epsilon$, and for any other $k$ we can substitute it into Eqn 10 and verify that it satisfies the equation:

$$1-(1-\epsilon)^{k+1} = \epsilon + (1-\epsilon)\left[1-(1-\epsilon)^k\right] = \epsilon + 1 - \epsilon - (1-\epsilon)^{k+1}, \tag{12}$$

the two sides being equal. Eqn 11 describes an exponentially saturating curve, which means that the likelihood of extinction simply compounds by the same factor across trophic levels. Given the linear assumption on the response to prey loss $w$, this is an intuitive result.

We now compare the expected number of total vs. direct extinctions in the network. Direct extinctions are those one would expect without taking the food web context into account. Let us say there are $m$ trophic levels in total. Then, since there are $n$ species per level, and every species has an extinction probability of $\epsilon$ (since the network is ignored), the expected number of direct extinctions $D$ is simply

$$D = mn\epsilon. \tag{13}$$

In turn, the expected number of *total* extinctions $T$ is what one would expect given the network structure. Adding up all the extinction probabilities of the species using Eqn 11, we get

$$T = \sum_{k=1}^{m} np_k = n\sum_{k=1}^{m}\left[1-(1-\epsilon)^k\right] = mn - n(1-\epsilon)\frac{1-(1-\epsilon)^m}{1-(1-\epsilon)}. \tag{14}$$

The Excess Extinction $E$ of a food web is given by the ratio of $T$ and $D$:

$$E = \frac{T}{D} = \frac{mn - n(1-\epsilon)\frac{1-(1-\epsilon)^m}{1-(1-\epsilon)}}{mn\epsilon}. \tag{15}$$

Simplifying, this reduces to

$$E = \frac{(m+1)\epsilon + (1-\epsilon)^{m+1} - 1}{m\epsilon^2}. \tag{16}$$

The Fragility $F$ of a food web is given by the ratio of $T-D$ and $S-D$, where $S = mn$ is the total number of species in the food web:

$$F = \frac{T-D}{S-D} = \frac{T/D-1}{S/D-1} = \frac{E-1}{S/D-1}. \tag{17}$$

Since $S = mn$ and $D = mn\epsilon$, we have $S/D = 1/\epsilon$ and the above simplifies to

$$F = \frac{E-1}{1/\epsilon - 1} = \frac{\epsilon}{1-\epsilon}(E-1). \tag{18}$$

Applying this to Eqn 16 and simplifying, we obtain the Fragility:

$$F = \frac{\epsilon}{1-\epsilon}\left(\frac{(m+1)\epsilon + (1-\epsilon)^{m+1} - 1}{m\epsilon^2} - 1\right) = \frac{m\epsilon + (1-\epsilon)^m - 1}{m\epsilon}. \tag{19}$$

#### 3 Excess Extinction and Fragility in the convex nonlinear case

We now calculate Excess Extinction $E$ and Fragility $F$ under the assumption of $w(i/n) = (i/n)^2$. With this choice, Eqn 6 reads

$$p_C = \epsilon + (1-\epsilon)\sum_{i=0}^{n}\left(\frac{i}{n}\right)^2\binom{n}{i}p_R^i(1-p_R)^{n-i}. \tag{20}$$

The sum is related to the second moment of the binomial distribution, and can thus be written as

$$\sum_{i=0}^{n}\left(\frac{i}{n}\right)^2\binom{n}{i}p_R^i(1-p_R)^{n-i} = p_R\left(p_R + \frac{1-p_R}{n}\right). \tag{21}$$

Substituting this into Eqn 20, we get

$$p_C = \epsilon + (1-\epsilon)p_R\left(p_R + \frac{1-p_R}{n}\right). \tag{22}$$

Compared with the analogous Eqn 9 in the linear case of $w(i/n) = i/n$, there is an unpleasant corollary to this equation. Namely, its right hand side depends explicitly on the number of resource species $n$. This in turn implies that consumers at the same trophic level no longer have the same associated marginal extinction probability: the number of connections matter. Even worse, it is not just the raw connectance value, but the actual structure of connections that will determine the extinction probabilities of the species at any given trophic level. Consequently, one cannot write a generic recursive equation like Eqn 10 in which all species are assumed to have the same marginal extinction probabilities $p_k$ at trophic level $k$.

There is one way out of this predicament. Looking at Eqn 22, we see that the dependence on $n$ is not very strong as long as $n$ is not too small. Thus, if we assume that every consumer is connected to at least a handful of resource species, then the $n$-dependence of the marginal extinction probabilities will be weak at best. Formally, we write the equation in the $n \to \infty$ limit where the $(1-p_R)/n$ term disappears:

$$p_C = \epsilon + (1-\epsilon)p_R^2. \tag{23}$$

Now once again we have an equation that does not depend on $n$ and thus does not depend on connectance. Furthermore, this also implies that the marginal extinction probabilities $p_k$ at any trophic level $k$ will be equal. Eqn 23 can therefore be written as a recursive equation:

$$p_{k+1} = \epsilon + (1-\epsilon)p_k^2, \tag{24}$$

with initial condition $p_1 = \epsilon$.

Unfortunately, Eqn 24 is still too complicated to be solved explicitly. But one can employ an approximation, which is based on the fact that the dynamical behavior of this equation is very smooth: it simply converges to a fixed point without any oscillations (permanent or damped). To show this, let us obtain its fixed points $p^*$:

$$p^* = \epsilon + (1-\epsilon)p^{*2}, \tag{25}$$

which has the two roots

$$p_1^* = 1, \qquad p_2^* = \frac{\epsilon}{1-\epsilon}. \tag{26}$$

Let us look at their stability, by calculating the Jacobian and evaluating it at these fixed points:

$$J = \frac{\partial p_{k+1}}{\partial p_k} = 2(1-\epsilon)p_k, \tag{27}$$

$$\lambda_1 = J|_{p_1^*} = 2(1-\epsilon), \tag{28}$$

$$\lambda_2 = J|_{p_2^*} = 2\epsilon. \tag{29}$$

In fact, $J = 2(1-\epsilon)p_k \geq 0$ for every $p_k \in [0,1]$ and not merely at the fixed points, so the map is monotonically increasing on the whole unit interval and oscillations are ruled out globally. In turn, $p_1^*$ is stable ($\lambda_1 < 1$) for $\epsilon > 1/2$ while $p_2^*$ is manifestly stable ($\lambda_2 < 1$) for $\epsilon < 1/2$. At $\epsilon = 1/2$, the two fixed points coincide and exchange stability in a transcritical bifurcation.

Since $p_k$ must be between 0 and 1, there is always one unique stable fixed point for any value of $\epsilon$. Starting from $p_1 = \epsilon$ (which lies below that fixed point), the monotonicity we just established means the trajectory increases towards it without ever leaving $[0,1]$. This means we can replace Eqn 24 with a corresponding differential equation, in which $k$ is allowed to vary continuously. Subtracting $p_k$ from both sides of Eqn 24 and employing the approximation $p_{k+1} - p_k \approx \mathrm{d}p(k)/\mathrm{d}k$, we get

$$\frac{\mathrm{d}p(k)}{\mathrm{d}k} = \epsilon - p(k) + (1-\epsilon)p(k)^2, \tag{30}$$

or, omitting the argument of $p$ for notational simplicity:

$$\frac{\mathrm{d}p}{\mathrm{d}k} = \epsilon - p + (1-\epsilon)p^2. \tag{31}$$

Writing Eqn 31 as $\mathrm{d}p/\mathrm{d}k = f(p)$, the recursion Eqn 24 is exactly $p_{k+1} = p_k + f(p_k)$, i.e., a forward Euler step of unit size applied to Eqn 31. The two therefore share the same fixed points, and their stability conditions correspond exactly, since the multiplier of the map is $\lambda = 1 + f'(p^*)$: the map is stable ($|\lambda| < 1$) precisely when $-2 < f'(p^*) < 0$, which is the stability condition for the differential equation whenever $f'(p^*) > -2$. Here $f'(p_1^*) = 1 - 2\epsilon$ and $f'(p_2^*) = 2\epsilon - 1$. Both are within that range, so no spurious stability change is introduced by the continuum approximation.

Eqn 31 can be integrated by separating variables:

$$\int \frac{\mathrm{d}p}{\epsilon - p + (1-\epsilon)p^2} = \int \mathrm{d}k. \tag{32}$$

The right-hand side evaluates simply to $k$. The left hand side integrates to

$$\int \frac{\mathrm{d}p}{\epsilon - p + (1-\epsilon)p^2} = \frac{\log(p-1) - \log(p - \epsilon - p\epsilon)}{1-2\epsilon} + K, \tag{33}$$

where $K$ is an integration constant. Its value is found using the fact that $p(k=1)=\epsilon$:

$$\frac{\log(\epsilon-1)-\log(-\epsilon^2)}{1-2\epsilon}+K=1, \tag{34}$$

from which

$$K=1-\frac{\log(\epsilon-1)-\log(-\epsilon^2)}{1-2\epsilon}. \tag{35}$$

So Eqn 32 evaluates to

$$\frac{\log(p-1)-\log(p-\epsilon-p\epsilon)}{1-2\epsilon}+1-\frac{\log(\epsilon-1)-\log(-\epsilon^2)}{1-2\epsilon}=k, \tag{36}$$

which can be solved for $p$ to give

$$p=\epsilon\frac{\mathrm{e}^{2\epsilon+k}(1-\epsilon)-\mathrm{e}^{2k\epsilon+1}\epsilon}{\mathrm{e}^{2\epsilon+k}(1-\epsilon)^2-\mathrm{e}^{2k\epsilon+1}\epsilon^2}. \tag{37}$$

We can now use this solution to calculate the $D$, $T$, $E$, and $F$. The expected number of direct extinctions $D$ is the number of species times $\epsilon$, like in Eqn 13:

$$D=mn\epsilon, \tag{38}$$

where $m$ is the number of trophic levels. The expected number of total extinctions $T$ will be approximated by integrating Eqn 37 across the trophic levels. Since the sum $\sum_{k=1}^{m} p_k$ assigns to each level $k$ a unit interval centered on $k$, the corresponding integral must run from $1/2$ to $m+1/2$ (equivalently, the sum is the midpoint-rule approximation of that integral):

$$T=n\sum_{k=1}^{m}p_k\approx n\int_{1/2}^{m+1/2}p(k)\,\mathrm{d}k=\int_{1/2}^{m+1/2}n\epsilon\frac{\mathrm{e}^{2\epsilon+k}(1-\epsilon)-\mathrm{e}^{2k\epsilon+1}\epsilon}{\mathrm{e}^{2\epsilon+k}(1-\epsilon)^2-\mathrm{e}^{2k\epsilon+1}\epsilon^2}\,\mathrm{d}k. \tag{39}$$

The primitive function $G(k)$ can be evaluated:

$$G(k)=\int n\epsilon\frac{\mathrm{e}^{2\epsilon+k}(1-\epsilon)-\mathrm{e}^{2k\epsilon+1}\epsilon}{\mathrm{e}^{2\epsilon+k}(1-\epsilon)^2-\mathrm{e}^{2k\epsilon+1}\epsilon^2}\,\mathrm{d}k=n\frac{k\epsilon-\log\left[(1-\epsilon)^2-\epsilon^2\mathrm{e}^{(k-1)(2\epsilon-1)}\right]}{1-\epsilon}, \tag{40}$$

and so the expected number of total extinctions is given by $T=G(m+1/2)-G(1/2)$. Excess Extinction $E$ is therefore

$$E=\frac{T}{D}=\frac{G(m+1/2)-G(1/2)}{D}, \tag{41}$$

or

$$E=\frac{m\epsilon+\log\left[\dfrac{(1-\epsilon)^2-\epsilon^2\mathrm{e}^{(1-2\epsilon)/2}}{(1-\epsilon)^2-\epsilon^2\mathrm{e}^{(m-1/2)(2\epsilon-1)}}\right]}{m\epsilon(1-\epsilon)}. \tag{42}$$

Fragility is then obtained from Eqn 18 as

$$F=\frac{\epsilon}{1-\epsilon}\left(\frac{m\epsilon+\log\left[\dfrac{(1-\epsilon)^2-\epsilon^2\mathrm{e}^{(1-2\epsilon)/2}}{(1-\epsilon)^2-\epsilon^2\mathrm{e}^{(m-1/2)(2\epsilon-1)}}\right]}{m\epsilon(1-\epsilon)}-1\right). \tag{43}$$

Eqn 42 approximates well the "true" solution obtained by explicitly iterating Eqn 24, for any number of trophic levels. The quality of approximation is shown in Figure 1 for various numbers of trophic levels.

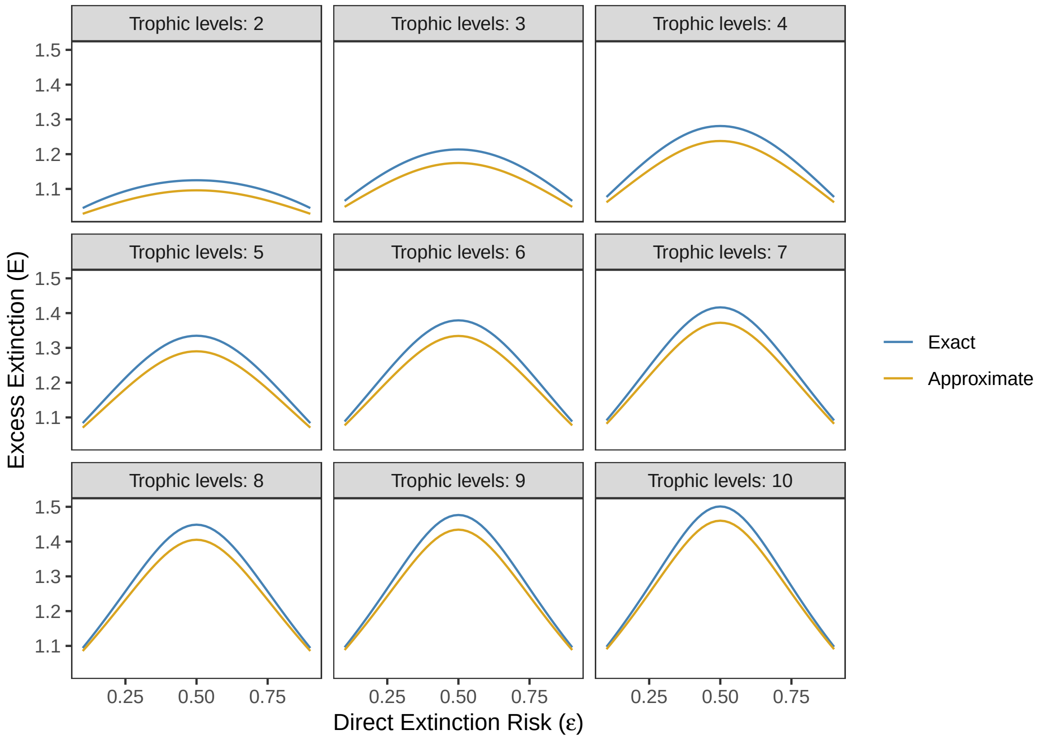


Figure 1: Excess Extinction (ordinate) for various direct extinction risks $\epsilon$ (abscissa) and number of trophic levels (panels), using both the exact values by numerically solving Eqn 24 and computing Excess Extinction (blue) and the approximation of Eqn 42 (yellow). The approximation is qualitatively correct for every number of levels shown, and captures the non-monotonic behavior of Excess Extinction as a function of $\epsilon$.

#### 4 Excess Extinction and Fragility in the concave nonlinear case

Now we use the same approach to calculate Excess Extinction $E$ and Fragility $F$ for the the choice $w(i/n) = 1 - (1 - (i/n))^2 = 2(i/n) - (i/n)^2$. Eqn 6 now reads

$$p_C = \epsilon + (1-\epsilon) \sum_{i=0}^{n} \left[ 2\left(\frac{i}{n}\right) - \left(\frac{i}{n}\right)^2 \right] \binom{n}{i} p_R^i (1-p_R)^{n-i}. \tag{44}$$

By using the first and second moments of the binomial distribution, we get

$$p_C = \epsilon + 2(1-\epsilon)p_R - (1-\epsilon)p_R \left( p_R + \frac{1-p_R}{n} \right). \tag{45}$$

Again writing the equation in the $n \to \infty$ limit, we can eliminate the $(1-p_R)/n$ term:

$$p_C = \epsilon + (1-\epsilon)(2p_R - p_R^2). \tag{46}$$

This implies that the marginal probabilities of extinction $p_k$ at any trophic level $k$ will be equal, and so we can write Eqn 46 as a recursive equation:

$$p_{k+1} = \epsilon + (1-\epsilon)(2p_k - p_k^2), \tag{47}$$

with initial condition $p_1 = \epsilon$.

As it happens, Eqn 47 can be solved explicitly:

$$p_k = 1 - (1-\epsilon)^{2^k - 1}. \tag{48}$$

One could now use this to obtain $E$ and $F$, but doing so would involve sums over the above expression for $p_k$, which do not simplify out. To get a more explicit approximation, we rely on the same strategy as in Section 3: we solve a corresponding differential equation. To do so, first we find the fixed points of Eqn 47:

$$p^* = \epsilon + (1-\epsilon)(2p^* - p^{*2}), \tag{49}$$

which has the two roots

$$p_1^* = 1, \qquad p_2^* = \frac{\epsilon}{\epsilon - 1}. \tag{50}$$

Now observe that $p_2^* < 0$ for all $\epsilon \in (0,1)$, but we require that $p_2^* \geq 0$ since it represents a probability of extinction. Thus $p_1^*$ is the only valid fixed point that we consider. We calculate the Jacobian and evaluate it at $p_1^*$:

$$J = \frac{\partial p_{k+1}}{\partial p_k} = 2(1-\epsilon)(1-p_k), \tag{51}$$

$$\lambda_1 = J|_{p_1^*} = 0. \tag{52}$$

Thus $p_1^*$ is superstable ($\lambda_1 = 0$) for all $\epsilon \in (0,1)$. As in Section 3, $J = 2(1-\epsilon)(1-p_k) \geq 0$ on the whole of $[0,1]$, so the map is monotonically increasing there, and the trajectory starting from $p_1 = \epsilon$ rises to $p_1^*$ without oscillation and without leaving the unit interval. As in Section 3, this justifies converting the recursion into a differential equation via $p_{k+1} - p_k \approx \mathrm{d}p(k)/\mathrm{d}k$:

$$\frac{\mathrm{d}p}{\mathrm{d}k} = \epsilon + (1-2\epsilon)p - (1-\epsilon)p^2. \tag{53}$$

This differential equation can be integrated by separating variables:

$$\int \frac{\mathrm{d}p}{\epsilon + (1-2\epsilon)p - (1-\epsilon)p^2} = \int \mathrm{d}k. \tag{54}$$

The right hand side evaluates to $k$, and the left hand side integrates to

$$\int \frac{\mathrm{d}p}{\epsilon + (1-2\epsilon)p - (1-\epsilon)p^2} = \log((1-\epsilon)p+\epsilon) - \log(1-p) + K, \tag{55}$$

where $K$ is an integration constant. Using the fact that $p(k=1)=\epsilon$, we have

$$\log(2\epsilon - \epsilon^2) - \log(1-\epsilon) + K = 1, \tag{56}$$

and so

$$K = 1 + \log(1-\epsilon) - \log(2\epsilon - \epsilon^2). \tag{57}$$

So Eqn 54 evaluates to

$$\log((1-\epsilon)p+\epsilon) - \log(1-p) + 1 + \log(1-\epsilon) - \log(2\epsilon-\epsilon^2) = k, \tag{58}$$

which can be solved for $p$ to give

$$p = \frac{\mathrm{e}^{k-1}\epsilon(2-\epsilon) - \epsilon(1-\epsilon)}{\mathrm{e}^{k-1}\epsilon(2-\epsilon) + (1-\epsilon)^2}. \tag{59}$$

The expected number of direct extinctions follows Eqn 13 again: $D = mn\epsilon$, where $m$ is the number of trophic levels. We approximate the expected number of total extinctions $T$ by integrating Eqn 59 across the trophic levels, with the integration limits running from 1/2 to $m+1/2$ (for the same reason as in Eqn 39):

$$T = n\sum_{k=1}^{m} p_k \approx n\int_{1/2}^{m+1/2} p(k)\,\mathrm{d}k = \int_{1/2}^{m+1/2} n\frac{\mathrm{e}^{k-1}\epsilon(2-\epsilon) - \epsilon(1-\epsilon)}{\mathrm{e}^{k-1}\epsilon(2-\epsilon) + (1-\epsilon)^2}\,\mathrm{d}k. \tag{60}$$

The primitive function $G(k)$ can be evaluated:

$$G(k) = \int n\frac{\mathrm{e}^{k-1}\epsilon(2-\epsilon) - \epsilon(1-\epsilon)}{\mathrm{e}^{k-1}\epsilon(2-\epsilon) + (1-\epsilon)^2}\,\mathrm{d}k = \frac{n}{1-\epsilon}\log\left[\frac{\mathrm{e}^{k-1}\epsilon(2-\epsilon) + (1-\epsilon)^2}{\left(\mathrm{e}^{k-1}\epsilon(2-\epsilon)\right)^{\epsilon}}\right], \tag{61}$$

and so the expected number of total extinctions is given by $T = G(m+1/2) - G(1/2)$. The Excess Extinction $E$ is therefore

$$E = \frac{T}{D} = \frac{G(m+1/2) - G(1/2)}{D}, \tag{62}$$

or

$$E = \frac{\log\left[\frac{\mathrm{e}^{m-1/2}\epsilon(2-\epsilon) + (1-\epsilon)^2}{\mathrm{e}^{-1/2}\epsilon(2-\epsilon) + (1-\epsilon)^2}\right] - m\epsilon}{m\epsilon(1-\epsilon)}. \tag{63}$$

Using Eqn 18 to obtain the Fragility:

$$F = \frac{\epsilon}{1-\epsilon}\left(\frac{\log\left[\frac{\mathrm{e}^{m-1/2}\epsilon(2-\epsilon) + (1-\epsilon)^2}{\mathrm{e}^{-1/2}\epsilon(2-\epsilon) + (1-\epsilon)^2}\right] - m\epsilon}{m\epsilon(1-\epsilon)} - 1\right). \tag{64}$$

Eqn 63 approximates well the true solution obtained by summing Eqn 48 across the trophic levels. The quality of approximation is shown in Figure 2 for various numbers of trophic levels.

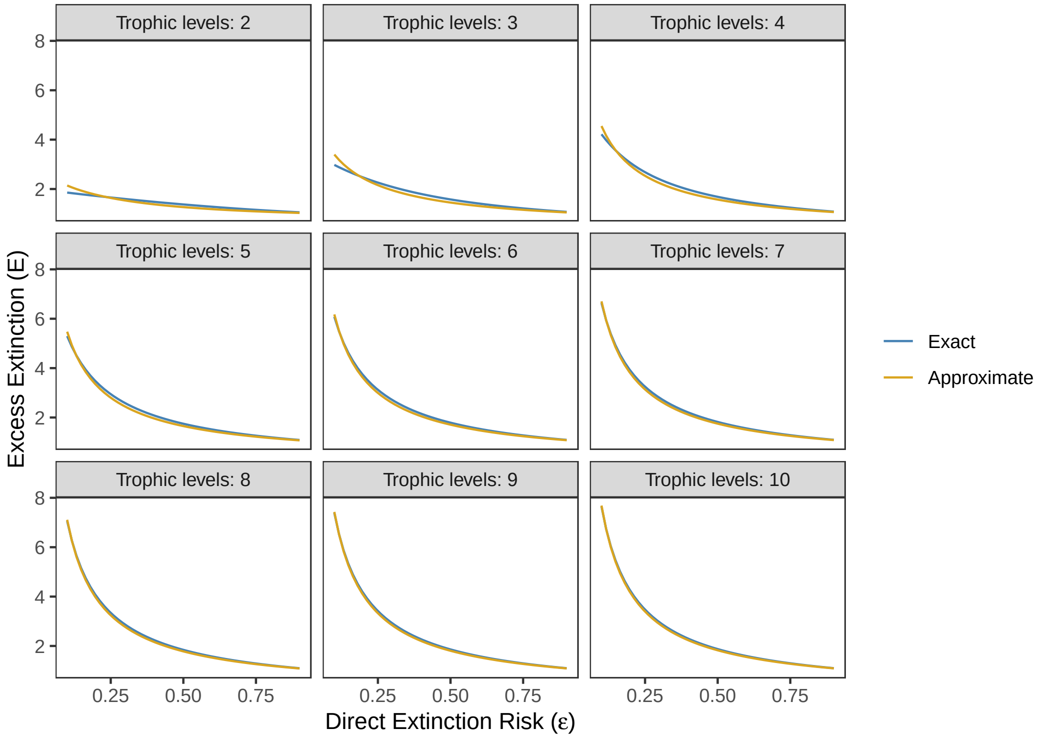


Figure 2: Excess Extinction (ordinate) for various direct extinction risks $\epsilon$ (abscissa) and number of trophic levels (panels), using both the exact values by summing Eqn 48 and computing Excess Extinction (blue) and the approximation of Eqn 63 (yellow).
