## Supplementary material for "Maximum trophic level predicts food webs’ susceptibility to coextinctions": SuppInfo_text_figures_tables

*Niche model food web generation*

For a given food web size $S$ and target connectance $C$, the niche model assigns each species in the food web a "niche value" and probabilistically determines feeding interactions based on the relative ranking of species' niche values. Since the interaction assigning mechanism of the niche model is stochastic in nature, the realized connectance of a food web generated by the niche model will deviate slightly from the desired target connectance. This deviation is usually negligible, and as described below, we ensured that the realized connectances of food webs generated by the niche model were sufficiently close to the predefined target connectance values before proceeding with the analysis.

While these two models of synthetic food webs require different parameters, we can generate comparable stratified and niche food webs by setting the niche model's number of species to $S = m \cdot n$ and its target connectance to $C = (q \cdot n(m - 1)) / (m(m \cdot n - 1))$, where $m$ is the number of trophic levels in the stratified web and $n$ is the number of species per trophic level in the stratified web. For example, a stratified food web with $m = 3$, $n = 4$, and $q = 0.5$ will have the same size and expected connectance as a niche model food web with $S = 12$ and $C \approx 0.121$ (Figure S1). Differences in the outcome of extinction scenario simulations on these two food webs can then be attributed to differences in the link-assigning mechanisms of the two models and the resulting differences in food web structure.

During the food web generation process, we rejected candidate food webs that contained isolated species (those with no resources and no consumers) and trophically identical species (two species with identical sets of resources and consumers). We also rejected candidate food webs that were not connected (i.e., the food web could be partitioned into distinct unconnected components) and those with realized connectance that deviated from its target connectance by more than 5%. Finally, we removed all cannibalistic links (self-loops) and trophic cycles (directed cycles), since the Bayesian network approach requires acyclic graphs as input.

*Interaction of direct extinction risk and number of trophic levels in stratified food webs*

When the response function $w(f)$ is linear or concave, the importance of trophic level for Excess Extinction is magnified at lower values of direct extinction risk $\varepsilon$, while it has virtually no effect when $\varepsilon$ is close to 1 (Figures 2a and 2b). The effect of trophic level on Fragility is also dependent on the magnitude of direct extinction risk: $F$ increases nearly linearly with $m$ when $\varepsilon$ is small, but the relationship becomes increasingly concave as $\varepsilon$ increases (Figures 2d and 2e).

*Role of omnivorous interactions in predictive performance of stratified food webs*

Since one of the key structural differences between stratified food webs and more complex food webs (i.e., empirical food webs and those generated by the niche model) is the presence of omnivorous interactions, we investigated whether an index of omnivory could explain the relative accuracy of predictions made by stratified food webs: if an empirical food web contains very few omnivorous interactions and is nearly "stratified," then it is reasonable to expect that $E$ and $F$ are closely aligned with the predicted outcome made by corresponding stratified food webs. We defined an omnivory index as the mean standard deviation of the trophic level of a species' resources, averaged across all species within a food web. This is a non-negative real number, with stratified food webs having an omnivory index of 0. Next, we calculated a deviation score as the ratio of $E$ obtained for the empirical food web and $E$ predicted by its corresponding stratified food webs.

We find a weak correlation between the omnivory index and deviation score in our 12 empirical food webs ($\rho$ = 0.3131; Figure S8). This suggests that the improved performance of predictions made by stratified food webs is not due to a relative lack of omnivorous interactions in empirical food webs. In fact, the deviation in extinction outcomes between empirical food webs and their corresponding niche model food webs seems largely due to the fact that the number of trophic levels in empirical food webs is much lower than the number of trophic levels in corresponding food webs generated by the niche model (Figure S9).

*Role of functional form for response to resource loss*

The non-monotonic relationship between $E$ and $\varepsilon$ in simulations with a convex response function can be explained intuitively: at low levels of direct extinction risk $\varepsilon$, few species will go extinct as a direct result of the threat, and $D$ is small. Because consumers are less sensitive to an initial loss of prey, there are few coextinctions – $T$ is only slightly larger than $D$, and so $E$ is only marginally greater than 1. At high levels of direct extinction risk $\varepsilon$, most species will go extinct as a direct result of the threat, and $D$ is large. The remaining surviving species are very likely to become coextinct because they have lost the majority of their resources. $D$ and $T$ are both near their maximal value, the total number of species in the community, and thus $E$ is still close to 1. Only at moderate values of $\varepsilon$ are there few enough direct extinctions – yet sufficiently many to allow coextinctions to propagate through the food web – to yield a relatively high value of $T$ relative to $D$, thereby increasing $E$.

*Role of threat type*

We also investigate the sensitivity of our findings to the type of threat that species face. In addition to a deterministic threat, we conducted extinction simulations using a stochastic threat and a basal-only threat. We find that extinction outcomes for both $E$ and $F$ under a stochastic threat very closely approximate those under a deterministic threat (top and middle rows of Figures S10-S15).

Under a basal-only threat, however, there are large differences in both $E$ and $F$ compared to a deterministic or stochastic threat. When the response function is linear, $E$ increases linearly with $m$ and $\varepsilon$ has no effect (Figure S10g). Conversely, $F$ increases with $\varepsilon$ and $m$ has virtually no effect (Figure S11g). When the response function is concave, $E$ and $F$ show qualitatively similar behavior under a basal-only threat as they do under a deterministic or stochastic threat (middle columns of Figures S10 and S11). However, the magnitude of $E$ is significantly larger for all values of $\varepsilon$ and $m$ under a basal-only threat (Figure S10h). Finally, when the response function is convex, $E$ increases with both $m$ and $\varepsilon$, and the non-monotonic relationship between $E$ and $\varepsilon$ is no longer observed (Figure S10i). $F$ still increases with $\varepsilon$, but under a basal-only threat it decreases with $m$ (Figure S11i).

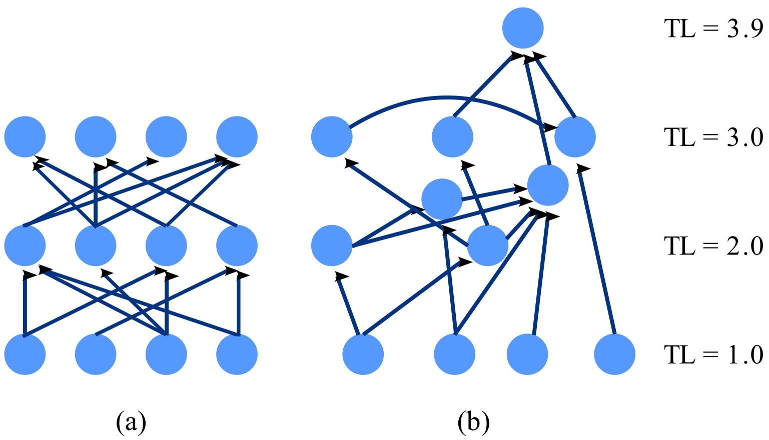


**Figure S1. Comparison of food web models.** A stratified food web (a) with $m = 3$ discrete trophic levels, $n = 4$ species per trophic level, and interaction probability $q = 0.5$. A niche model food web (b) with $S = 12$ species and connectance $C \approx 0.121$. These two food webs, while generated by different models, contain the same number of species and have approximately the same connectance. Trophic Level (TL) denotes the prey-averaged trophic level of each species in the two food webs. Note that while the niche model food web allows for omnivory (feeding at multiple trophic levels), the stratified food web only allows for feeding between adjacent trophic levels. Furthermore, species in the niche model food web can have non-integer trophic level.

| Name | Acronym | Reference | Ecosystem | Location | Species | Links | Connectance | Max TL |
|---|---|---|---|---|---|---|---|---|
| Mount Batten | EMB | Mendonca et al., 2018 | Rocky intertidal | South coast, England | 45 | 289 | 0.146 | 3.07 |
| Wembury | EWB | Mendonca et al., 2018 | Rocky intertidal | South coast, England | 45 | 346 | 0.175 | 3.43 |
| Porto da Cruz | MPC | Mendonca et al., 2018 | Rocky intertidal | Madeira Island, Portugal | 55 | 364 | 0.123 | 4.17 |
| Canico | MRM | Mendonca et al., 2018 | Rocky intertidal | Madeira Island, Portugal | 39 | 171 | 0.115 | 4.00 |
| Cabo Raso | PCR | Mendonca et al., 2018 | Rocky intertidal | West coast, Portugal | 86 | 777 | 0.106 | 4.77 |
| Raio Verde | PRV | Mendonca et al., 2018 | Rocky intertidal | West coast, Portugal | 92 | 748 | 0.089 | 4.54 |
| Chesapeake Bay | CHB | Baird and Ulanowicz, 1989 | Coastal salt marsh | Chesapeake Bay, USA | 31 | 67 | 0.072 | 3.90 |
| Bahia Falsa de San Quintin | BSQ | Hechinger et al., 2011 | Coastal salt marsh | Baja California, Mexico | 78 | 509 | 0.085 | 4.18 |
| Carpinteria Salt Marsh | CSM | Hechinger et al., 2011 | Coastal salt marsh | California, USA | 92 | 757 | 0.090 | 4.37 |
| Estero de Punta Banda | EPB | Hechinger et al., 2011 | Coastal salt marsh | Baja California, Mexico | 105 | 1090 | 0.100 | 4.42 |
| Little Rock Lake | LRL | Martinez, 1991 | Freshwater lake | Wisconsin, USA | 95 | 1088 | 0.122 | 4.44 |
| Saint Martin | STM | Goldwasser and Roughgarden, 1993 | Terrestrial | Saint Martin Island | 48 | 214 | 0.095 | 4.72 |

**Table S1. Description of empirical food webs used for analysis.** Max TL is the highest prey-averaged trophic level attained by any species in the food web.

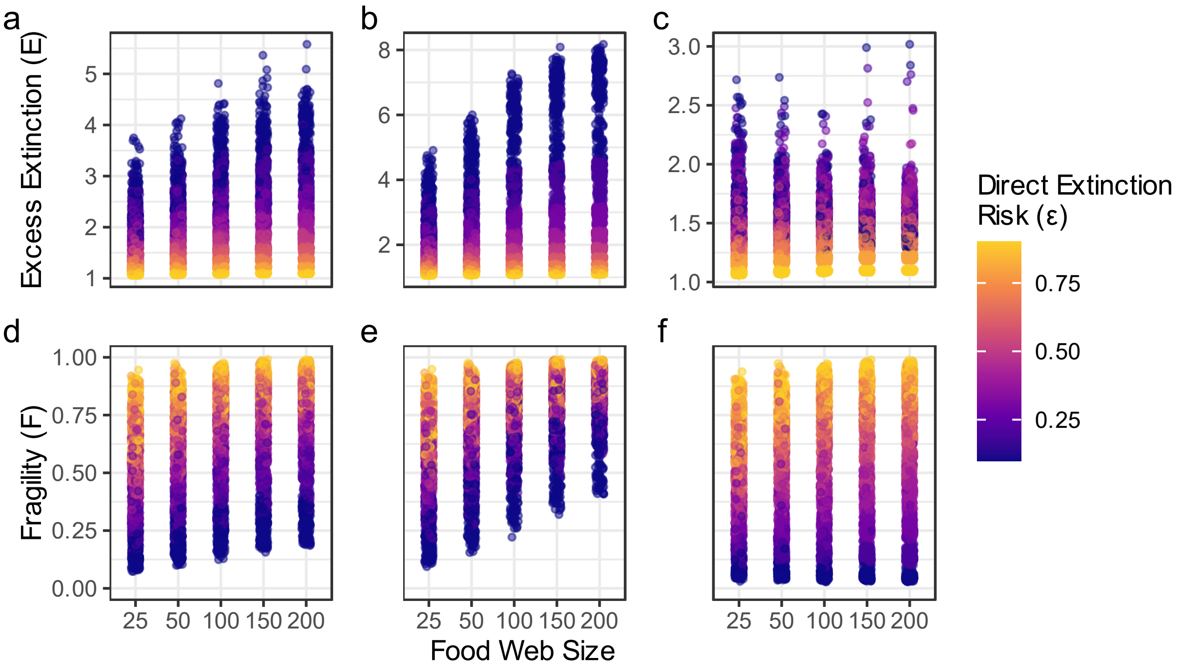


**Figure S2. Excess Extinction and Fragility as a function of food web size in niche model food webs.** Excess Extinction (top row) and Fragility (bottom row) as a function of niche model food web size with a deterministic threat. Each point describes the outcome of an extinction simulation (horizontal jitter added to reduce overplotting) when the response function $w(f)$ is linear (left column), concave (middle column), and convex (right column). Point color indicates ε, the direct extinction risk.

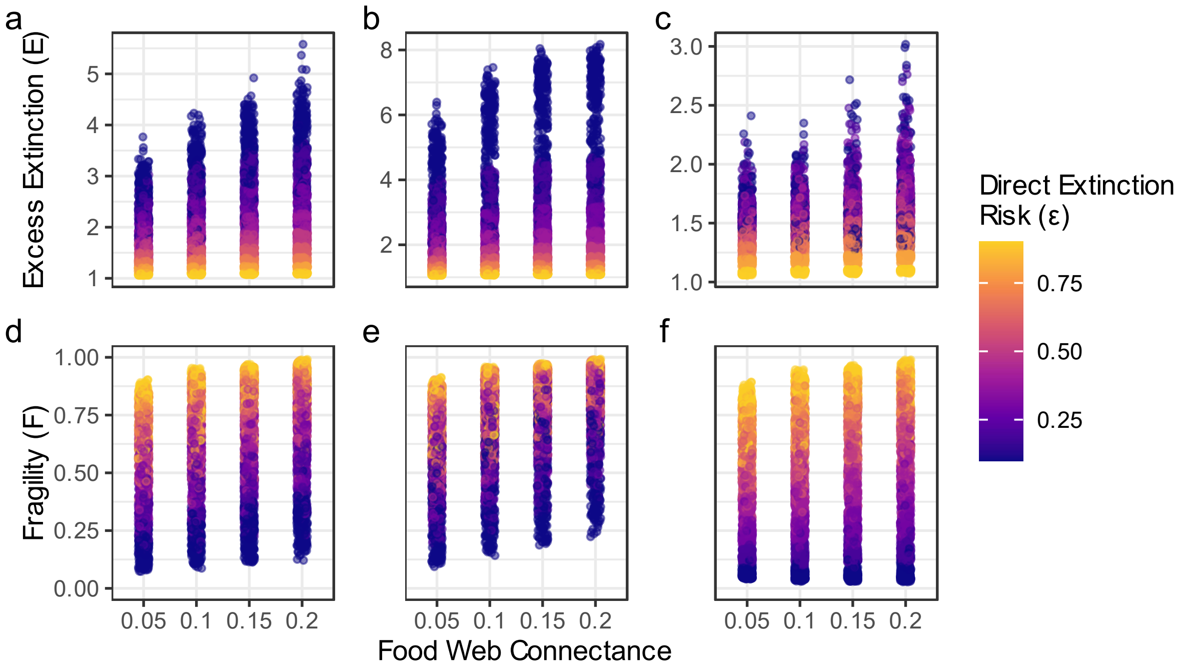


**Figure S3. Excess Extinction and Fragility as a function of food web connectance in niche model food webs.** Excess Extinction (top row) and Fragility (bottom row) as a function of niche model food web connectance with a deterministic threat. Each point describes the outcome of an extinction simulation (horizontal jitter added to reduce overplotting) when the response function *w*(*f*) is linear (left column), concave (middle column), and convex (right column). Point color indicates ε, the direct extinction risk.

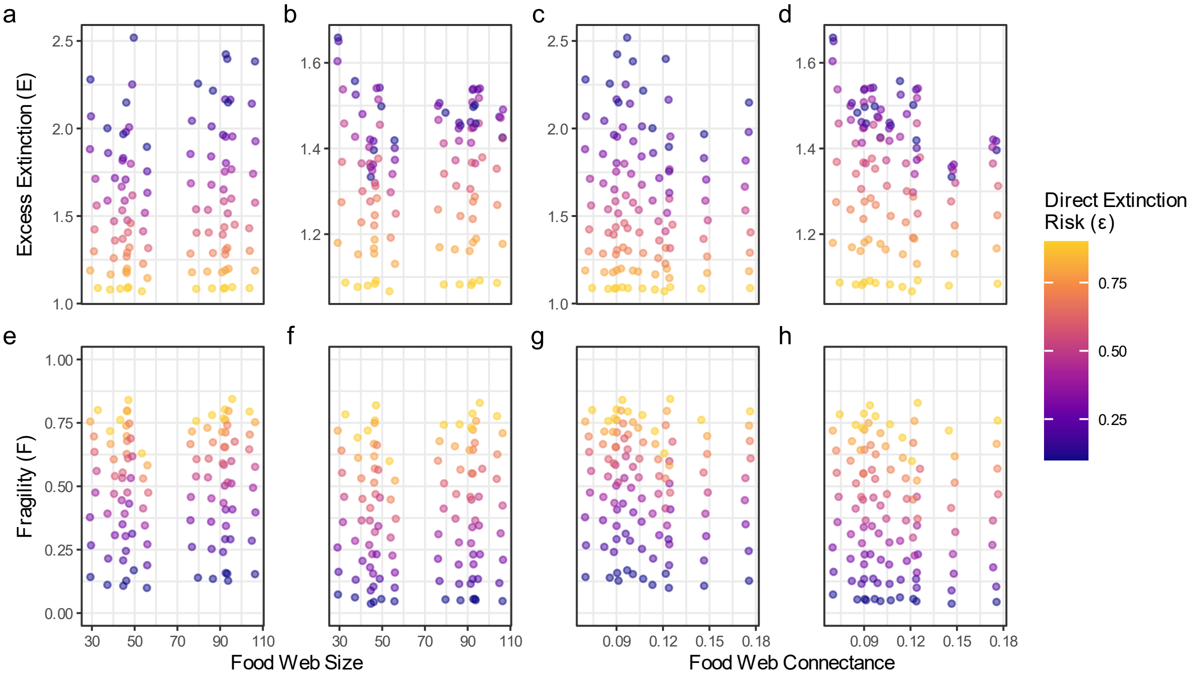


**Figure S4. Excess Extinction and Fragility as a function of food web size and connectance in empirical food webs.** Excess Extinction (top row) and Fragility (bottom row) as a function of food web size and connectance with a deterministic threat. Each point describes the outcome of an extinction simulation (horizontal jitter added to avoid overplotting) when the response function $w(f)$ is linear (a, c, e, and g) and convex (b, d, f, and h). Point color indicates ε, the direct extinction risk to each species. We find that food web size $S$ ranged from 31 to 105 species in our set of empirical food webs, while connectance $C$ ranged from 0.072 to 0.175 (n = 12).

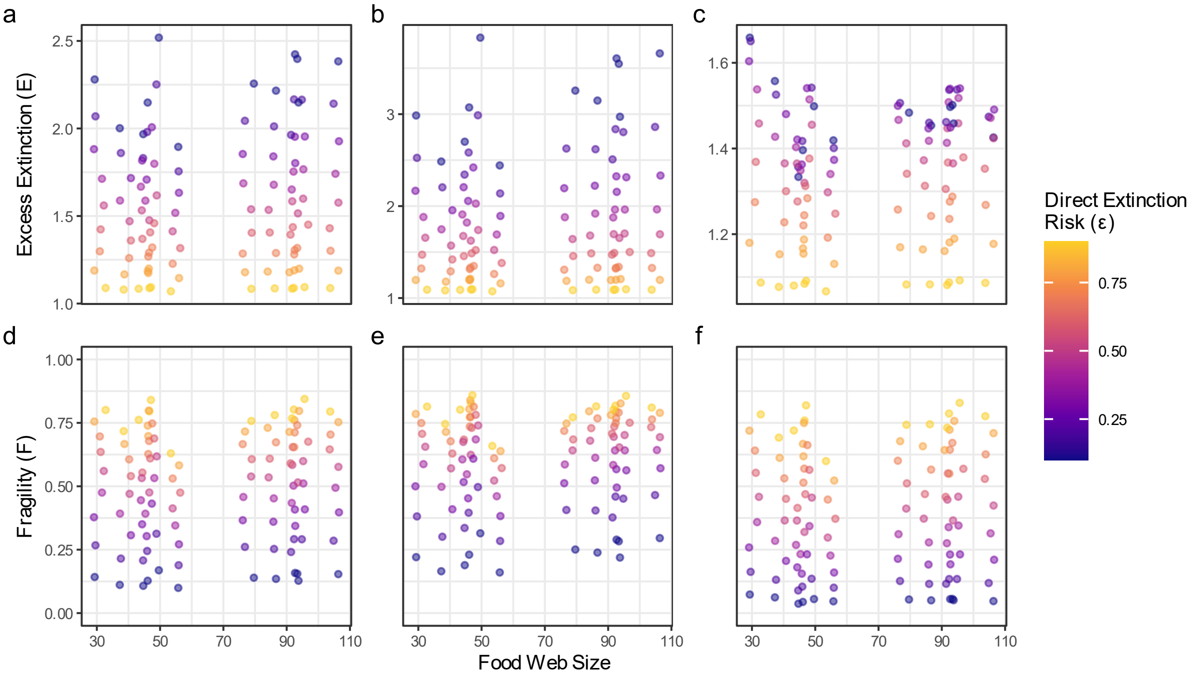


**Figure S5. Excess Extinction and Fragility as a function of food web size in empirical food webs.** Excess Extinction (top row) and Fragility (bottom row) as a function of empirical food web size with a deterministic threat. Each point describes the outcome of an extinction simulation (horizontal jitter added to reduce overplotting) when the response function $w(f)$ is linear (left column), concave (middle column), and convex (right column). Point color indicates ε, the direct extinction risk.

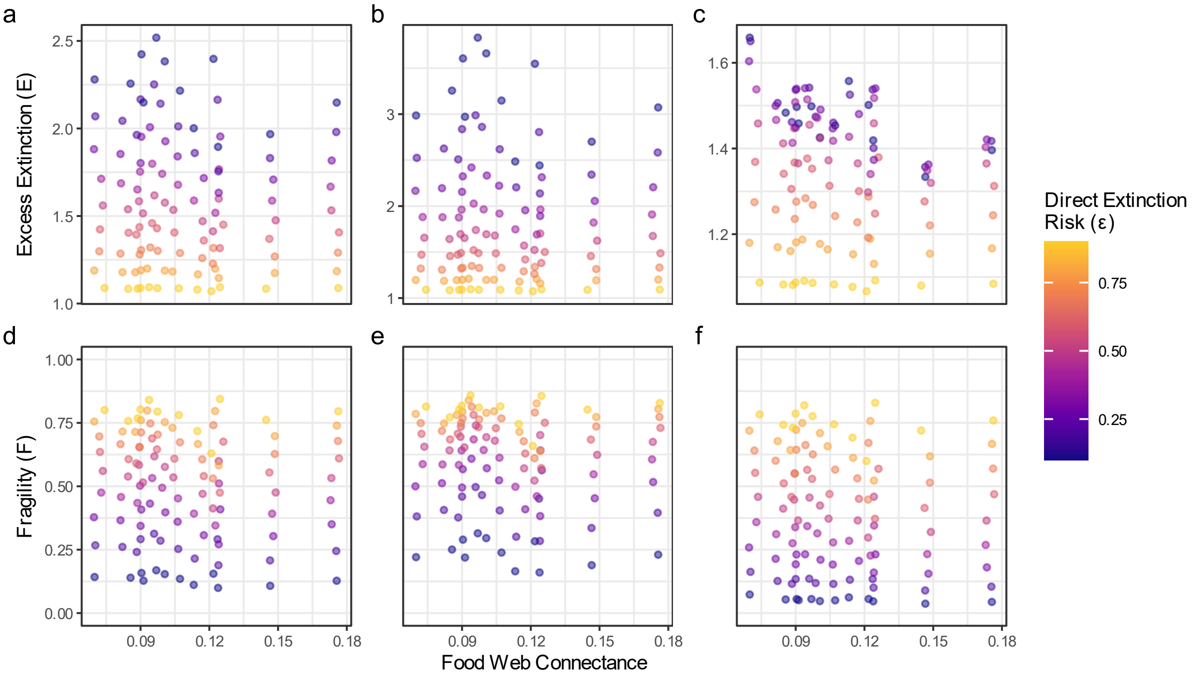


**Figure S6. Excess Extinction and Fragility as a function of food web connectance in empirical food webs.** Excess Extinction (top row) and Fragility (bottom row) as a function of empirical food web connectance with a deterministic threat. Each point describes the outcome of an extinction simulation (horizontal jitter added to reduce overplotting) when the response function *w*(*f*) is linear (left column), concave (middle column), and convex (right column). Point color indicates ε, the direct extinction risk.

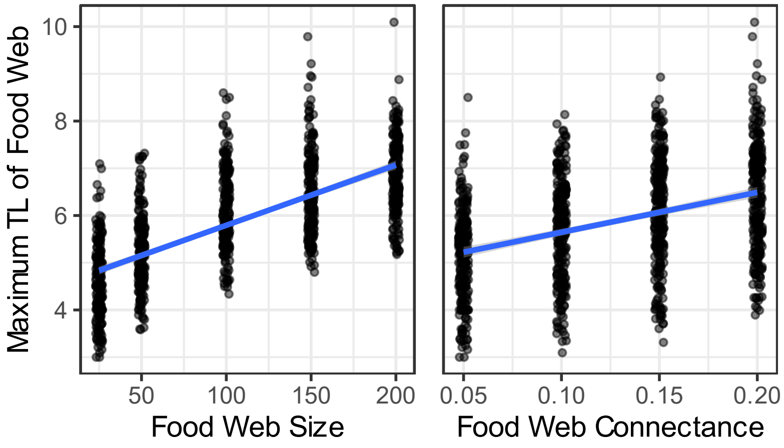


**Figure S7. Maximum trophic level in niche model food webs as a function of food web structure.** Each point describes the maximum trophic level, size (left panel), and connectance (right panel) of a food web generated by the niche model (horizontal jitter added to avoid overplotting). Blue lines indicate estimated linear relationships between maximum trophic level and food web structure.

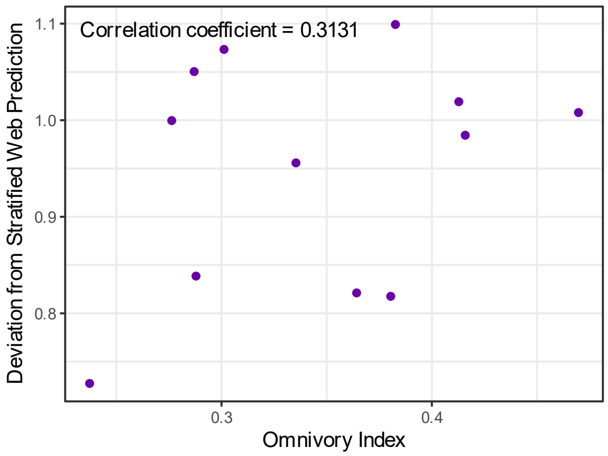


**Figure S8. Omnivory index of empirical food webs.** We measured the extent of omnivorous interactions present within each empirical food web (abscissa), as well as its deviation from the value of $E$ predicted by a corresponding stratified food web (ordinate). There is no significant correlation between food web omnivory index and accuracy of prediction generated by a corresponding stratified food web model parameterized by the maximum trophic level of the empirical food web.

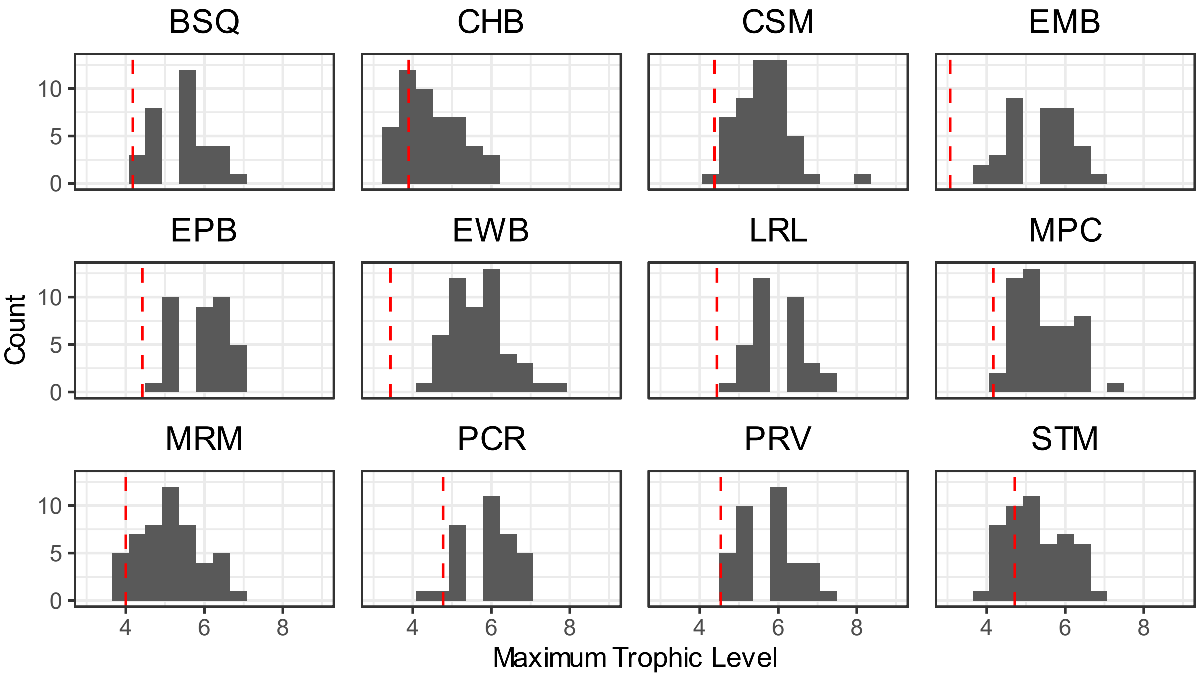


**Figure S9. Comparison of maximum trophic level in empirical food webs and corresponding niche model food webs.** We measured the maximum trophic level of each empirical food web (red dotted line), as well as distribution of maximum trophic levels for generated niche model food webs corresponding to each empirical food web (gray bars). In almost all cases, the maximum trophic level in the empirical food web is significantly lower than predicted by corresponding niche model food webs.

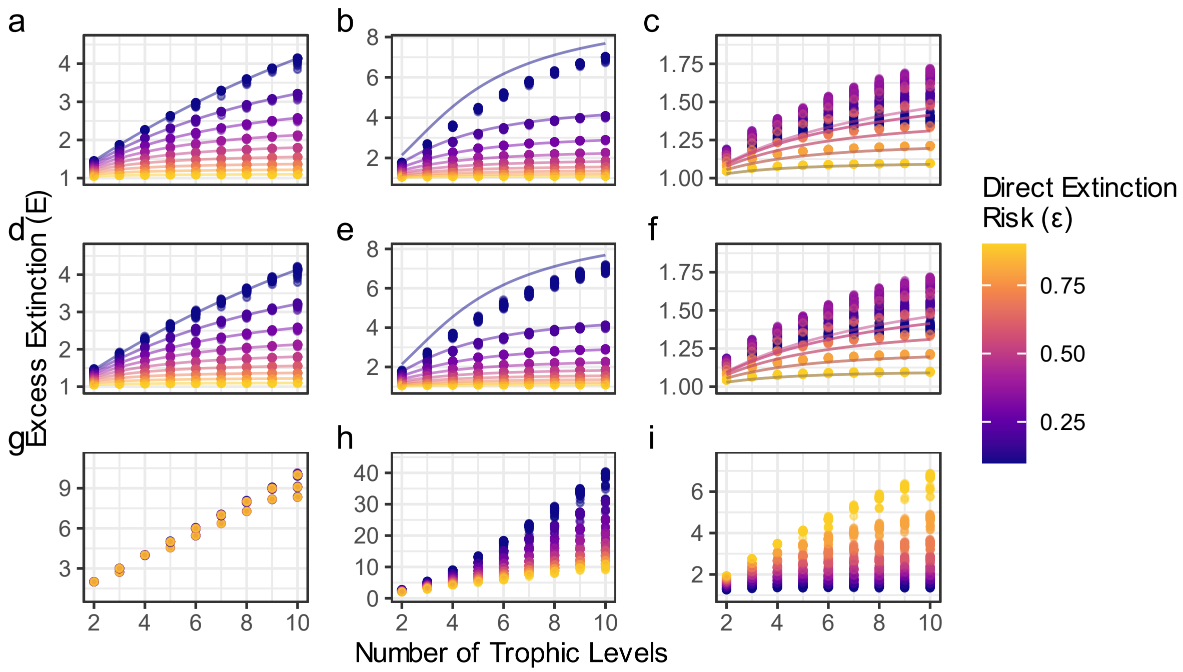


**Figure S10. Excess Extinction as a function of maximum trophic level in stratified food webs.** Each point describes the outcome of an extinction simulation (horizontal jitter added to avoid overplotting) when the response function $w(f)$ is linear (left column), concave (middle column), and convex (right column). Rows describe extinction simulation outcomes under a deterministic threat (top row), a stochastic threat (middle row), and a basal-only threat (bottom row). For the linear response function case, each line graphs $E$ as calculated by Equation 2. For the concave and convex cases, each line graphs approximations of $E$ as calculated by equations in Supporting Information. Note that only a subset of lines is shown in the convex case (c and f) because the equation for $E$ is symmetric about $\varepsilon = 0.5$, so some lines are covered by others (for example, the graph of $E$ is the same for $\varepsilon = 0.3$ and $\varepsilon = 0.7$). Point color indicates $\varepsilon$, the direct extinction risk to each species.

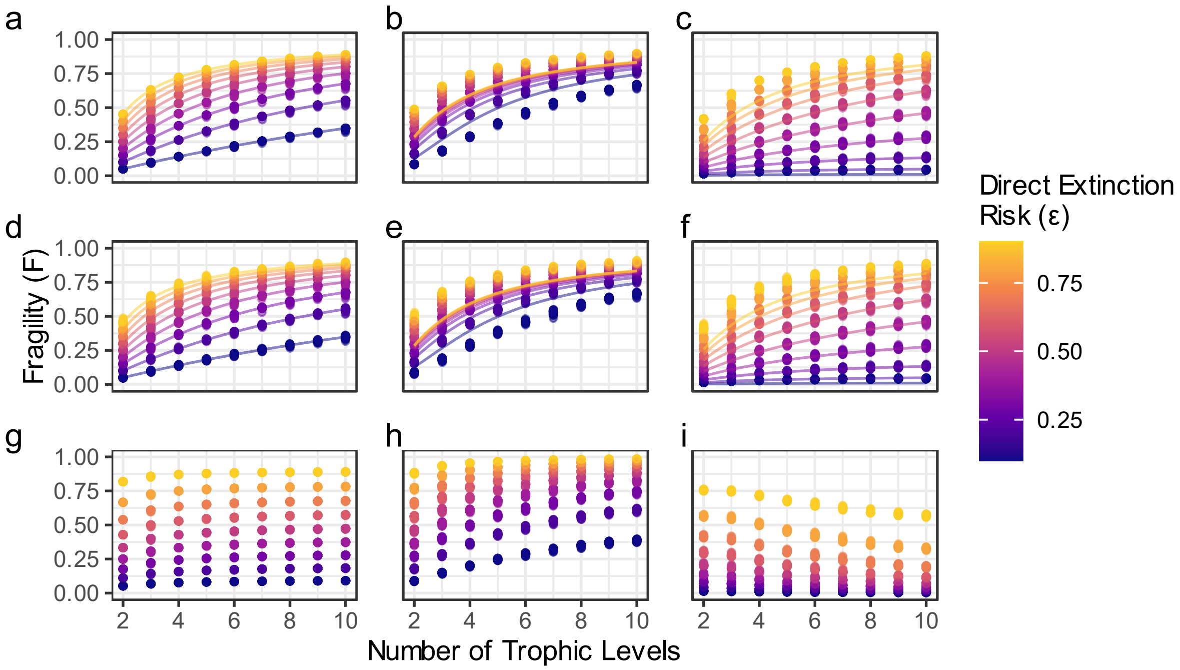


**Figure S11. Fragility as a function of maximum trophic level in stratified food webs.** Each point describes the outcome of an extinction simulation (horizontal jitter added to avoid overplotting) when the response function $w(f)$ is linear (left column), concave (middle column), and convex (right column). Rows describe extinction simulation outcomes under a deterministic threat (top row), a stochastic threat (middle row), and a basal-only threat (bottom row). For the linear response function case, each line graphs $F$ as calculated by Equation 3. For the concave and convex cases, each line graphs approximations of $F$ as calculated by equations in Supporting Information. Point color indicates ε, the direct extinction risk to each species.

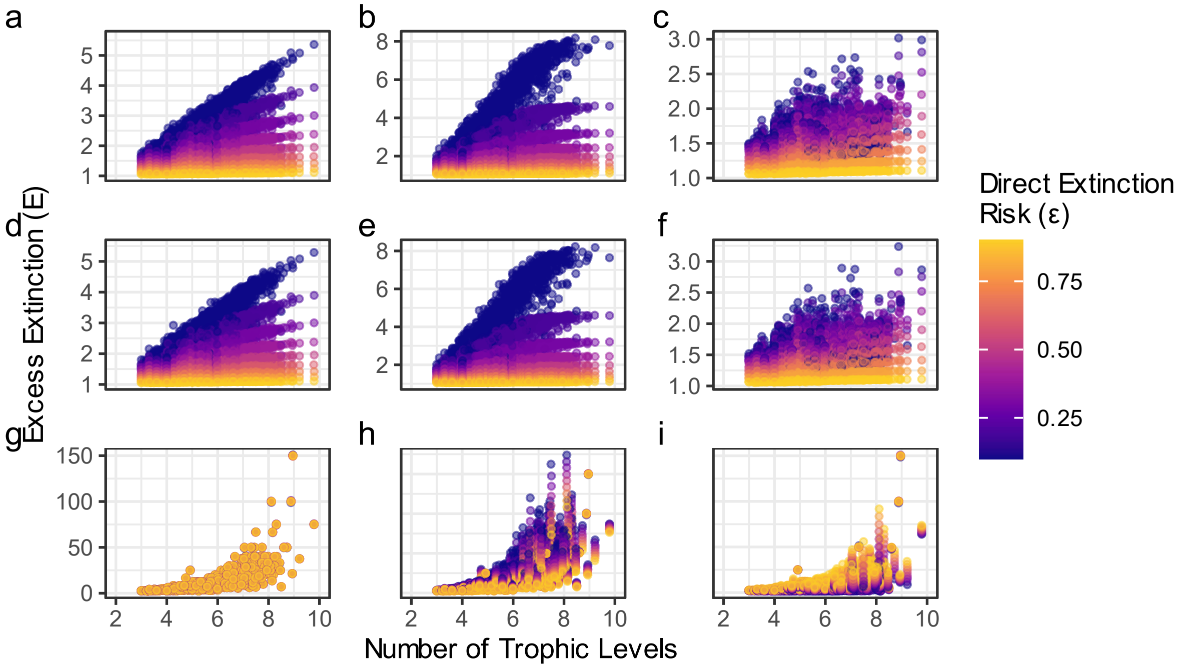


**Figure S12. Excess Extinction as a function of maximum trophic level in niche model food webs.** Each point describes the outcome of an extinction simulation (horizontal jitter added to avoid overplotting) when the response function $w(f)$ is linear (left column), concave (middle column), and convex (right column). Rows describe extinction simulation outcomes under a deterministic threat (top row), a stochastic threat (middle row), and a basal-only threat (bottom row). Point color indicates ε, the direct extinction risk to each species.

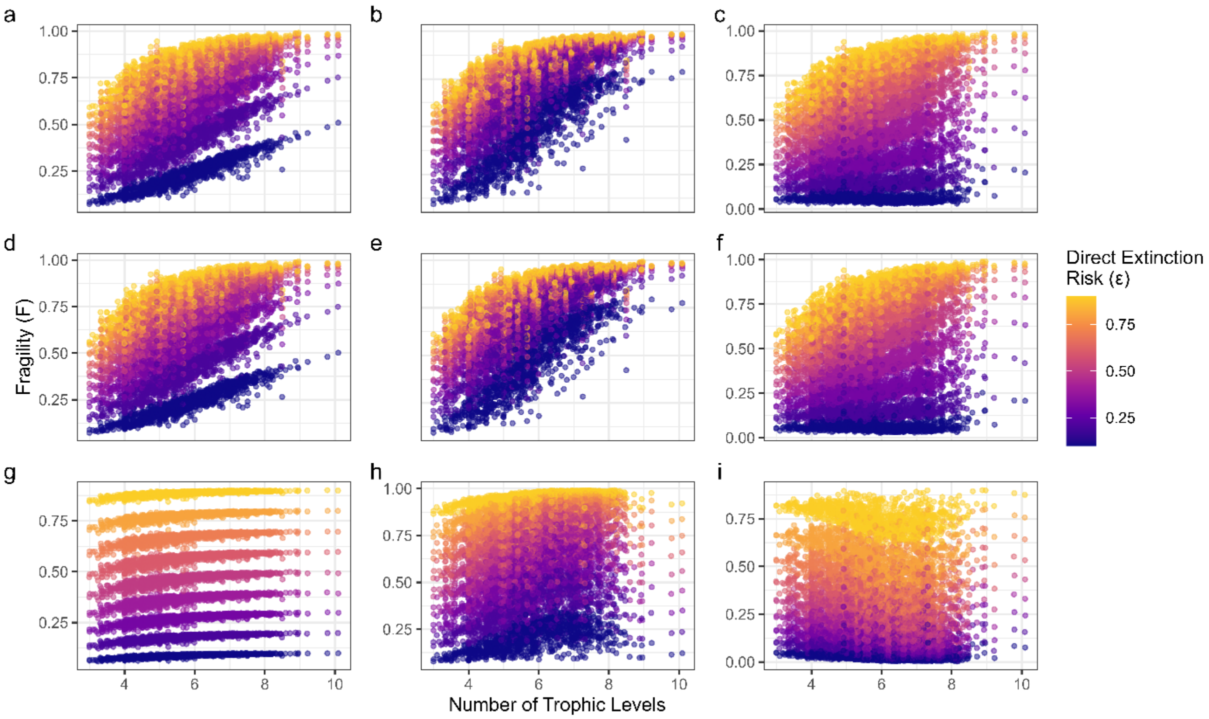


**Figure S13. Fragility as a function of maximum trophic level in niche model food webs.** Each point describes the outcome of an extinction simulation (horizontal jitter added to avoid overplotting) when the response function $w(f)$ is linear (left column), concave (middle column), and convex (right column). Rows describe extinction simulation outcomes under a deterministic threat (top row), a stochastic threat (middle row), and a basal-only threat (bottom row). Point color indicates ε, the direct extinction risk to each species.

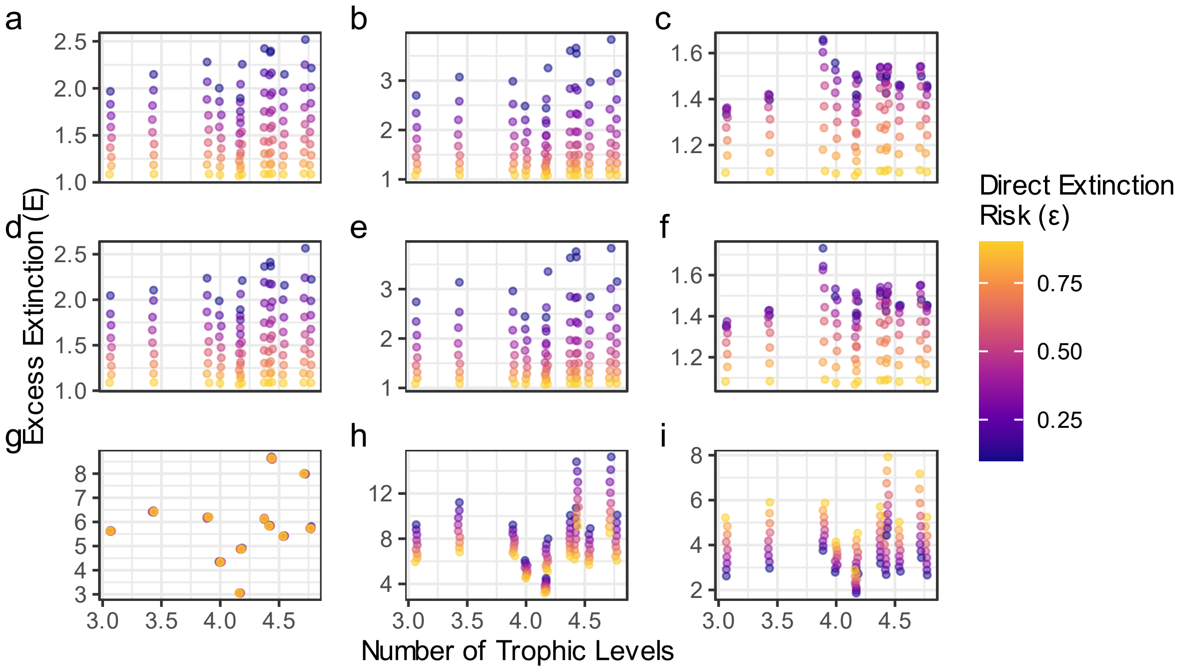


**Figure S14. Excess Extinction as a function of maximum trophic level in empirical food webs.** Each point describes the outcome of an extinction simulation (horizontal jitter added to avoid overplotting) when the response function *w*(*f*) is linear (left column), concave (middle column), and convex (right column). Rows describe extinction simulation outcomes under a deterministic threat (top row), a stochastic threat (middle row), and a basal-only threat (bottom row). Point color indicates ε, the direct extinction risk to each species.

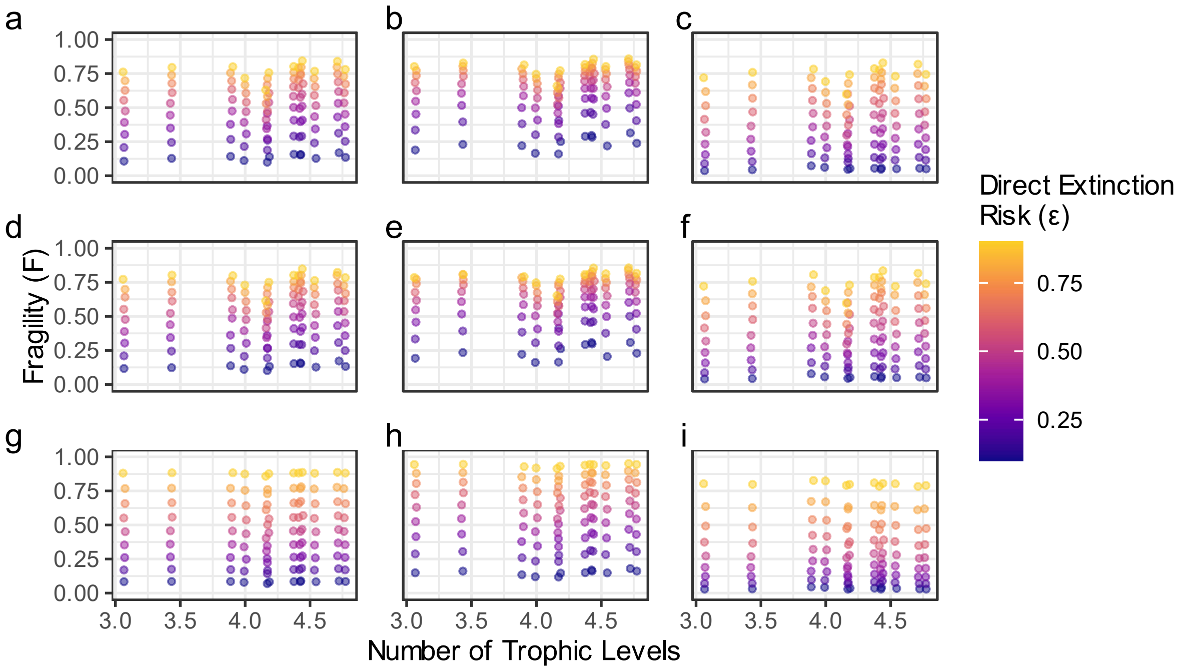


**Figure S15. Fragility as a function of maximum trophic level in empirical food webs.** Each point describes the outcome of an extinction simulation (horizontal jitter added to avoid overplotting) when the response function $w(f)$ is linear (left column), concave (middle column), and convex (right column). Rows describe extinction simulation outcomes under a deterministic threat (top row), a stochastic threat (middle row), and a basal-only threat (bottom row). Point color indicates ε, the direct extinction risk to each species.

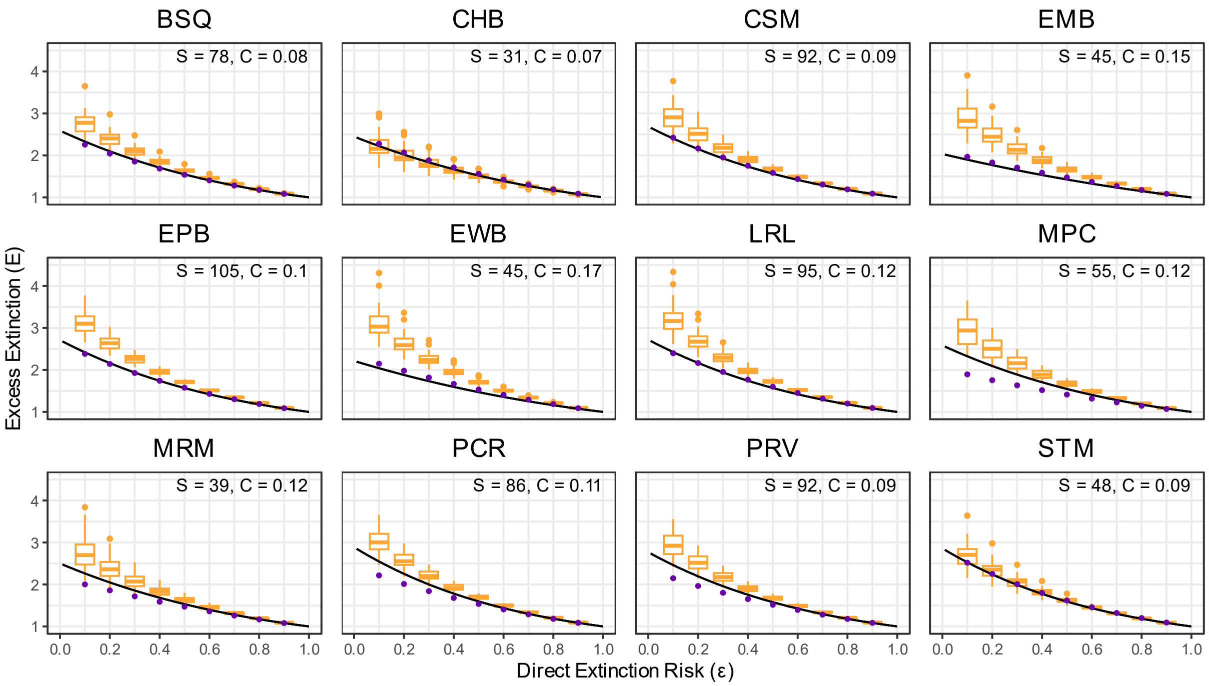


**Figure S16. Extinction simulation outcomes for empirical food webs compared to corresponding model food webs.** Extinction simulation outcomes using a linear response function *w*(*f*) and a deterministic threat are shown for all empirical food webs and their corresponding model food webs. Purple points indicate *E* for outcomes on empirical food webs. Golden box plots indicate the range of simulation outcomes for corresponding niche model food webs parameterized by the size and connectance of each empirical food web. Black lines graph *E* as calculated by Equation 2, parameterized by the maximum trophic level of each empirical food web.
